# The Sponges of the Carmel Pinnacles Marine Protected Area

**DOI:** 10.1101/2022.11.02.514922

**Authors:** Thomas L. Turner, Steve Lonhart

## Abstract

California’s network of marine protected areas was created to protect the diversity and abundance of native marine life, but the status of some taxa is very poorly known. Here we describe the sponges (phylum Porifera) from the Carmel Pinnacles State Marine Reserve, as assessed by a SCUBA-based survey in shallow waters. Of the 29 sponge species documented, 12 (41%) of them were previously unknown. Using a combination of underwater photography, DNA sequencing, and morphological taxonomy, we greatly improve our understanding of the status and distribution of previously described species and formally describe the new species as *Hymedesmia promina* sp. nov.*, Phorbas nebulosus* sp. nov.*, Clathria unoriginalis* sp. nov., *Clathria rumsena* sp. nov.*, Megaciella sanctuarium* sp. nov.*, Mycale lobos* sp. nov.*, Xestospongia ursa* sp. nov.*, Haliclona melissae* sp. nov.*, Halichondria loma* sp. nov.*, Hymeniacidon fusiformis* sp. nov.*, Scopalina carmela* sp. nov., and *Obruta collector* gen. nov., sp. nov. An additional species, *Lissodendoryx topsenti* (de Laubenfels 1930), is moved to *Hemimycale,* and *H. polyboletus* comb. nov., nom. nov. is created due to preoccupation by *H. topsenti* (Burton, 1929). Several of the new species appear to be rare and/or have very restricted distributions, as they were not found at comparative survey sites outside of Carmel Bay. These results illustrate the potential of qualitative presence/absence systematic surveys of understudied taxa to discover and document substantial novel diversity.

## Introduction

Central California is well-known for its rugged coastline and scenic beauty, where local geologic, oceanographic, and biogeographic features combine to create a mosaic of unique habitats and biodiversity hotspots (Blanchette *et al*. 2008; Valentine 1966). As a transitional zone between the warm-temperate waters south of Point Conception (34.5° N latitude) and the cold-temperate waters north of Point Arena (39° N latitude), Central California has species from both regions (Lonhart *et al*. 2019; Zacherl *et al*. 2003). Recognition of these biological resources led to the establishment of Monterey Bay National Marine Sanctuary in 1992, which prohibits exploitation of oil, gas, and minerals within a large portion of this region. Further protections followed the passage of the California Marine Life Protection Act in 1999, which led to a network of state-designated marine protected areas (MPAs) along the entire coast of California. In Central California, most of these MPAs are nested within the federal marine sanctuary. One of these, the Carmel Pinnacles State Marine Reserve, encompasses 6.7 km^2^ surrounding a series of steep granite walls and pinnacles that rise to the surface from 144 m depth. Local currents and deep, clear, upwelled water support walls covered with filter feeding invertebrates and peaks topped by giant kelp and bull kelp sheltering schools of rockfishes. A variety of microhabitats and dramatic vertical relief make the site a popular diving destination.

MPAs were created to protect biodiversity, but the diversity of some taxa has scarcely been investigated. This lack of systematic inventory greatly limits our ability to assess conservation potential or success. Qualitative (presence/absence) surveys of understudied taxa provide an effective and efficient way to improve this understanding (Mikkelsen & Cracraft 2001). Sponges (phylum Porifera) are among the least inventoried macroscopic taxa in California. The Monterey Peninsula is the best studied region in California in terms of sponge diversity, but past work has focused on the intertidal zone and deep water, with very little collection by divers (Hartman 1975; de Laubenfels 1932). Sponges can provide diverse ecosystem services, from habitat formation to nutrient cycling (Bell 2008; de Goeij *et al*. 2013). The roles that sponges play in shallow-water ecosystems in California are poorly known, but with California’s kelp forests facing unprecedented rates of change, baseline information about the distribution and abundance of sponges is urgently needed. Towards this end, we used the National Marine Sanctuary vessel the R/V Tegula to do the first survey of sponges at the Carmel Pinnacles. We documented 29 sponge species; remarkably, 12 (41%) of them were previously unknown. We combined molecular phylogenies and morphological taxonomy to formally describe these new species and improve our understanding of previously described taxa. Half of the newly described species are thus far known only from locations in Carmel Bay, with four species known only from the Carmel Pinnacles SMR. The presence of so many rare and restricted species illustrates the value of qualitative surveys of understudied taxa, and motivates continued support of systematic inventories in the region.

## Methods

### Collections

Two 45-minute dives were made on the inner pinnacle at the Carmel Pinnacles State Marine Reserve. The first was on 8/10/21, at 36.55910, −121.96630, max depth 18 m; the second on 9/22/21, at 36.55852, −121.96820, max depth 24 m. A two-diver team comprised of the two authors made both dives. In order to maximize time to collect unknown sponges, sponges that can be tentatively identified from field photos were documented but not collected. Field-identifiability of sponges is based on the experience of Thomas Turner, who has collected over 1000 sponge samples in California since 2019. Collected samples are first identified based on field characteristics and macro photos, then these identifications are tested using spicules and/or DNA sequencing. The reliability of field-identification depends on the uniqueness and consistency of a species’ morphology, and also the number of individuals tested. Many species cannot be identified in the field because their morphology is too similar to other sympatric species, but many species can be identified, at least some of the time, because some individuals have characteristic combinations of color, shape, and surface patterning. The species indicated in table 1 as identified from field photos are all reliably identified based on current knowledge. This could change if future work discovers cryptic species with similar field morphology in the region, so these identifications should be considered tentative.

**Table 1.**
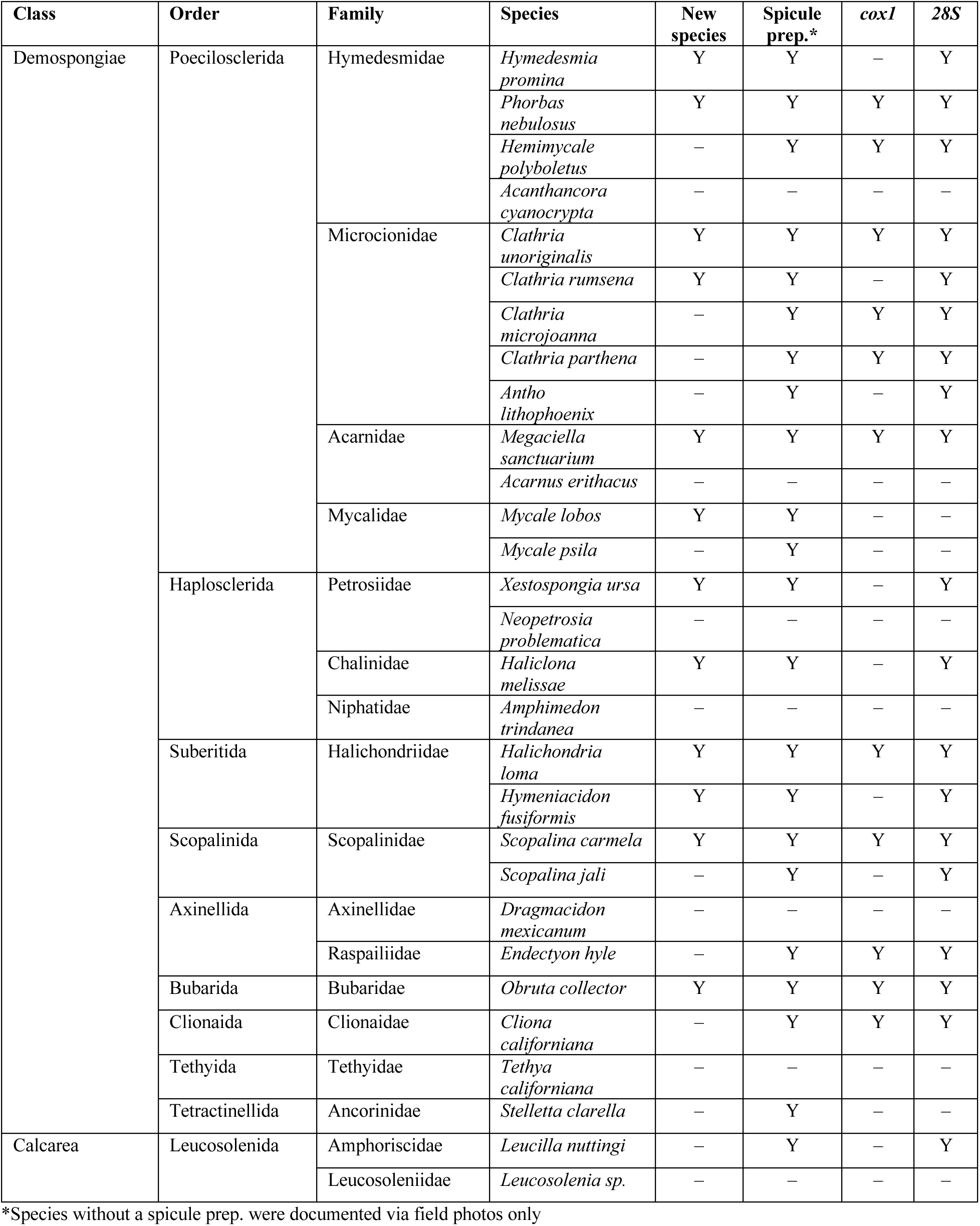
Taxonomic overview of collected samples and the standard of evidence supporting each identification (Y = yes, ‘–’ = no).

Samples of 28 sponges (from a total of 22 species) were hand-collected with a knife. Except for *Leucilla nuttingi*, only a portion of each sponge was sampled. An additional 54 samples are included as comparative material: 2 samples of *Xestospongia diprosopia* (de Laubenfels 1932) were loaned by the California Academy of Sciences in San Francisco, while the other 52 samples were drawn from recent collections made by Thomas Turner. These recent collections took place at 14 intertidal sites from Northern to Southern California, 23 marinas from Northern to Southern California, and via SCUBA at 14 subtidal sites around the Monterey Peninsula in Central California and 71 subtidal sites in Southern California.

Field-collected samples were placed in one-liter plastic bags with seawater, and these were kept on ice for up to 8 hours until sponges could be preserved in 95% EtOH. Each sample was assigned a collection number, preceded by TLT. Holotypes were vouchered with the California Academy of Sciences in San Francisco (voucher numbers with CASIZ). Pieces of most holotypes and many other samples were also vouchered at the Cheadle Center at the University of California, Santa Barbara (voucher numbers with IZC); paratypes and additional samples were vouchered with the Santa Barbara Museum of Natural History (voucher numbers with SBMNH). Several samples were collected during the “Los Angeles Urban Ocean Expedition 2019” bioblitz; these samples are archived at both the Natural History Museum of Los Angeles (voucher numbers with NHMLA) and the Florida Museum of Natural History (voucher numbers with BULA). Voucher numbers are listed for each species in the systematic section, but we also provide them in tabular format as supplementary data deposited at Data Dryad (https://doi.org/10.25349/D9ZS52). This supplementary table also includes additional metadata for each sample, such as Genbank numbers and collection locations.

### Morphology

Sponge subsamples were digested in bleach to examine spicules. Skeletal architecture was examined by hand cutting tissue sections and digesting them with a mixture of 97% Nuclei Lysis Solution (Promega; from the Wizard DNA isolation kit) and 3% 20 mg/ml Proteinase K (Promega). This digestion eliminates cellular material while leaving the spongin network intact. Sections of *Halichondria loma* sp. nov. were instead imaged by dehydrating in a 100% ethanol bath then clearing with Histoclear (National Diagnostics). Spicules and sections were imaged with a D3500 SLR camera (Nikon) with a NDPL-1 microscope adaptor (Amscope) attached to a compound trinocular microscope. Measurements were made on images using ImageJ (Schneider *et al*. 2012), after calculating the number of pixels per mm with a calibration slide. Spicule length was measured as the longest possible straight line from tip to tip, even when spicules were curved or bent. Spicule width was measured at the widest point, excluding adornments like swollen tyles. Scanning electron images were taken with a FEI Quanta400F Mk2 after coating spicules with 20 nm of carbon. Scale bars on figures are precise for microscopic images but often approximate for field photos. Spicule measurements in species descriptions are presented in the format minimum value – mean value – maximum value. Full distributions of spicule measurements are available as raw data at Data Dryad (https://doi.org/10.25349/D9ZS52).

### Genotyping

DNA extractions were performed with several different kits, all of which performed well (the Wizard Genomic DNA Purification kit (Promega), the Qiagen Blood & Tissue kit, and the Qiagen Powersoil kit). Three primer sets were used to sequence fragments of the *cox1* mitochondrial locus. Some samples were genotyped at a ∼1200 bp fragment using the following primers (LCO1490: 5-GGT CAA CAA ATC ATA AAG AYA TYG G-3’; COX1-R1: 5’-TGT TGR GGG AAA AAR GTT AAA TT-3’); these amplify the “Folmer” barcoding region and the “co1-ext” region used by some sponge barcoding projects (Rot *et al*. 2006). However, these primers failed to amplify a product for many samples. The Folmer region alone was amplified from these samples using the following primers: (LCO1490: 5’-GGT CAA CAA ATC ATA AAG AYA TYG G-3’; HCO2198: 5’-TAA ACT TCA GGG TGA CCA AAR AAY CA-3’) (Folmer *et al*. 1994). These primers nearly always amplified a product on freshly collected samples, but often fail to yield data due to preferentially amplifying DNA from contaminating algae, microbes, or marine invertebrates that are co-sampled with sponges. Neither primer set amplified a product from *Halichondria loma* sp. nov. due to primer mismatches, so custom primers were designed for this species: (halcox1-L: 5’-TGC TTG GCG ATG ATC ATT TA-3’; halcox1-R: 5’-ATG GCC CAA AGC ATA GGA GT-3’). These amplify an 875 bp fragment encompassing the folmer and cox-ext1 regions.

Three primer sets were used to amplify portions of the *28S* rDNA nuclear locus. Most species were sequenced across the D1-D5 region by combining two amplicons: the ∼800 bp D1-D2 region using primers Por28S-15F (5’-GCG AGA TCA CCY GCT GAA T-3’) and Por28S-878R (5’-CAC TCC TTG GTC CGT GTT TC-3’) and the ∼800 bp D3-D5 region using primers Por28S-830F (5’-CAT CCG ACC CGT CTT GAA-3’) and Por28S-1520R (5’-GCT AGT TGA TTC GGC AGG TG-3’) (Morrow *et al*. 2012). The D1-D2 primers failed to yield products for a small number of samples; for these, the C2-D2 region (covering ∼50% of the D1-D2 region) was amplified with C2 (5’-GAA AAG AAC TTT GRA RAG AGA GT-3’) and D2 (5’-TCC GTG TTT CAA GAC GGG-3’) (Chombard *et al*. 1998).

PCR was performed using a Biorad thermocycler (T100); the following conditions were used for the *cox1* locus: 95°C for 3 min, followed by 35 cycles of 94°C for 30 sec, 52°C for 30 sec, 72°C for 90 seconds, followed by 72°C for 5 minutes. For the halcox1 primers, this was modified to a 54°C annealing temperature. The *28S* C2-D2 region was amplified with the same conditions, except a 57°C annealing temperature and 60 second extension time; the *28S* D1-D2 and D3-D5 regions used a 53°C annealing temperature and 60 second extension time. PCR was performed in 50 μl reactions using the following recipe: 24 μl nuclease-free water, 10 μl 5x PCR buffer (Gotaq flexi, Promega), 8 μl MgCl, 1 μl 10mM dNTPs (Promega), 2.5 μl of each primer at 10 μM, 0.75 bovine serum albumin (10 mg/ml, final conc 0.15 mg/ml), 0.25 μl Taq (Gotaq flexi, Promega), 1 μl template. ExoSAP-IT (Applied Biosystems) was used to clean PCRs, which were then sequenced by Functional Biosciences using Big Dye V3.1 on ABI 3730xl instruments. Blastn was used to verify that the resulting traces were of sponge origin. All sequences have been deposited in GenBank under accession numbers OP526553-OP526599, OP534724-OP534744, OP535395-OP535398, and OP535457-OP535460; accession numbers are also shown in the phylogenies and listed in the supplementary table in Data Dryad (https://doi.org/10.25349/D9ZS52).

### Phylogenetic methods

Comparative sequences were collected from GenBank using a combination of blast and the NCBI taxonomy browser. For *cox1*, sequences were only included if they minimally encompassed the Folmer barcoding region. For *28S*, sequences were only included if they minimally encompassed the C2-D2 barcoding region. Phylogenies of each clade are not exhaustive, instead focusing on species closely related to local taxa and representative comparative groups. Sequence alignments were produced using MAFFT v.7 online (Katoh *et al*. 2017). Phylogenies were estimated with maximum likelihood using a GTR model in IQ-Tree (Nguyen *et al*. 2015; Trifinopoulos *et al*. 2016); the ultrafast bootstrap was used to measure node confidence (Hoang *et al*. 2018). Figures were produced by exporting IQ-Tree files to the Interactive Tree of Life webserver (Letunic & Bork 2019).

## Results

We documented 27 Demosponge species and two species of Calcarea in the Carmel Pinnacles SMR, as summarized in table 1. Diversity of higher taxa was substantial, with 10 orders and 17 families represented. Seven species were documented with field photos only (figure 1, table 1): these identifications should be considered tentative (see methods). In one case a sample was identified only to genus (*Leucosolenia* sp.): this is possibly *L. eleanor*, but the field identifiability of this group has not yet been tested in California.

**Figure 1.**
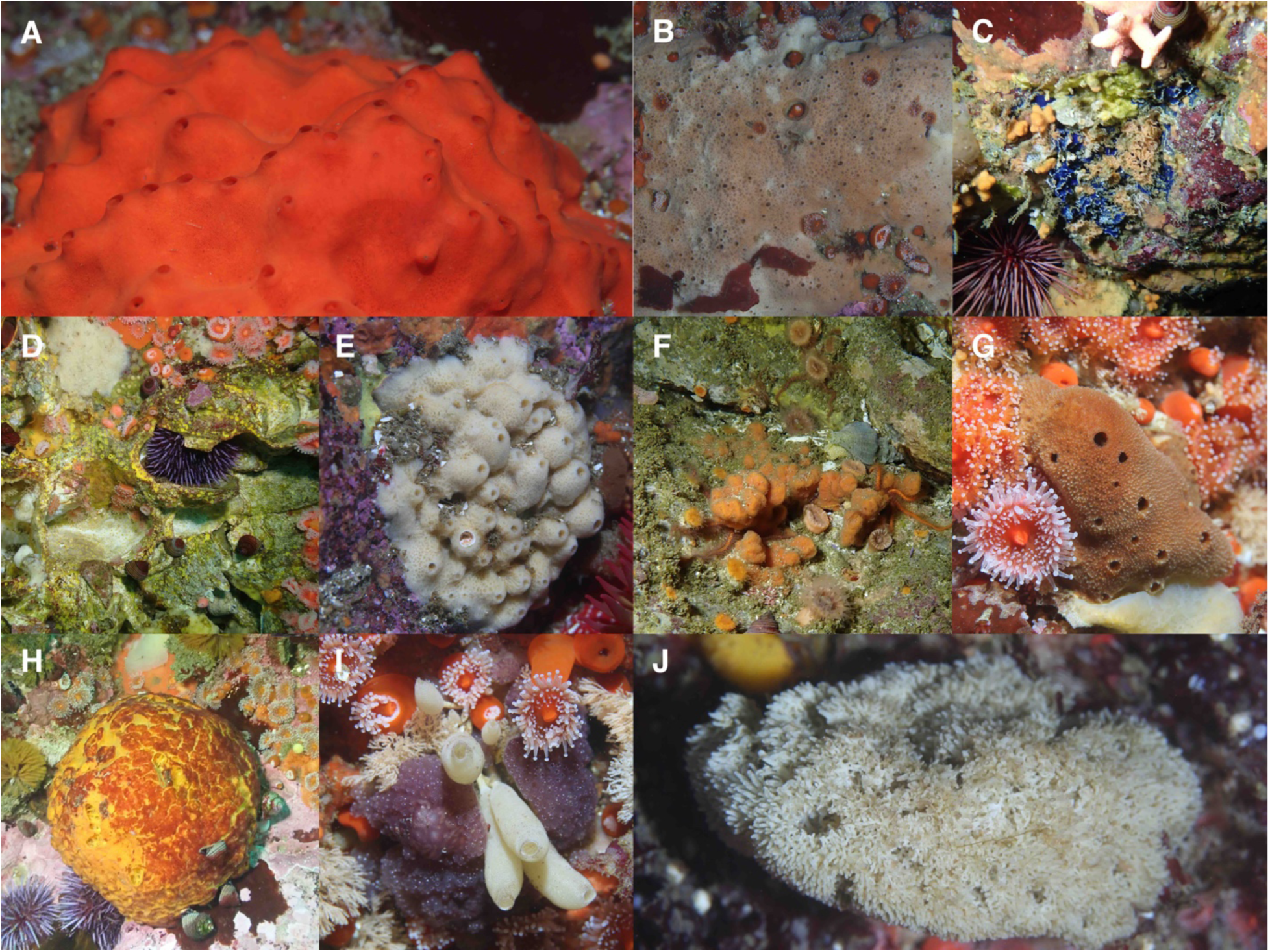
Assorted sponges from the Carmel Pinnacles. *Acarnus erithacus* (A), *Scopalina jali* (B), *Acanthancora cyanocrypta* (visible as a blue crust) (C), *Cliona californiana* (a boring species, visible as tiny yellow papillae) (D), *Neopetrosia problematica* (E), *Endectyon hyle* (F), *Amphimedon trindenea* (G), *Tethya californiana* (H), *Leucilla nuttingi* (I), *Leucosolenia* sp. (J). B, D, F, I are discussed in the systematics section below; other species were not collected, and are identified with field photos only.

In addition to the two collecting dives made at the Carmel Pinnacles, the authors also collected sponges at 13 other subtidal locations around the Monterey Peninsula. One of the authors (T. Turner) has also recently collected sponges at 71 subtidal locations in Southern California, 14 intertidal locations throughout California, and 23 marinas throughout California. Despite these extensive collections, four of the 12 new species have thus far only been found at the Carmel Pinnacles (*Hymedesmia promina* sp. nov.*, Megaciella sanctuarium* sp. nov.*, Hymeniacidon fusiformis* sp. nov., and *Scopalina carmela* sp. nov.). Two additional species were found at other locations in Carmel Bay, but not in Monterey Bay or any other location (*Clathria rumsena* sp. nov. and *Xestospongia ursa* sp. nov.); two species have been found only at sites in Carmel Bay plus San Miguel Island (*Phorbas nebulosus* sp. nov. and *Haliclona melissae* sp. nov.). The remaining 4 species were widespread, found throughout Central and Southern California (*Clathria unoriginalis* sp. nov., *Mycale lobos* sp. nov.*, Halichondria loma* sp. nov., and *Obruta collector* gen. nov., sp. nov.).

Collected samples were investigated both morphologically and genetically. DNA sequencing was focused on the two most common sponge barcoding loci (*cox1* and *28S*); amplification of both loci was attempted for all newly described species. Sequencing was successful for at least one locus for all new species except for *Mycale lobos*, where repeated and varied attempts were unsuccessful. DNA sequencing was also attempted for all previously described species because genetic data has proven critical in sponge systematics, and most California species have no previously available genetic data. Sequencing was unsuccessful for *Mycale psila* (de Laubenfels 1930) and *Stelletta clarella* (de Laubenfels 1930), but other species were successful and are included in updated phylogenies shown below. Comparative sequences were generated from additional individuals and comparative species in cases where these data increased confidence in species range determination and species delimitation, as described in the systematics section.

### Systematics

#### Order Poecilosclerida

**Figure 2.**
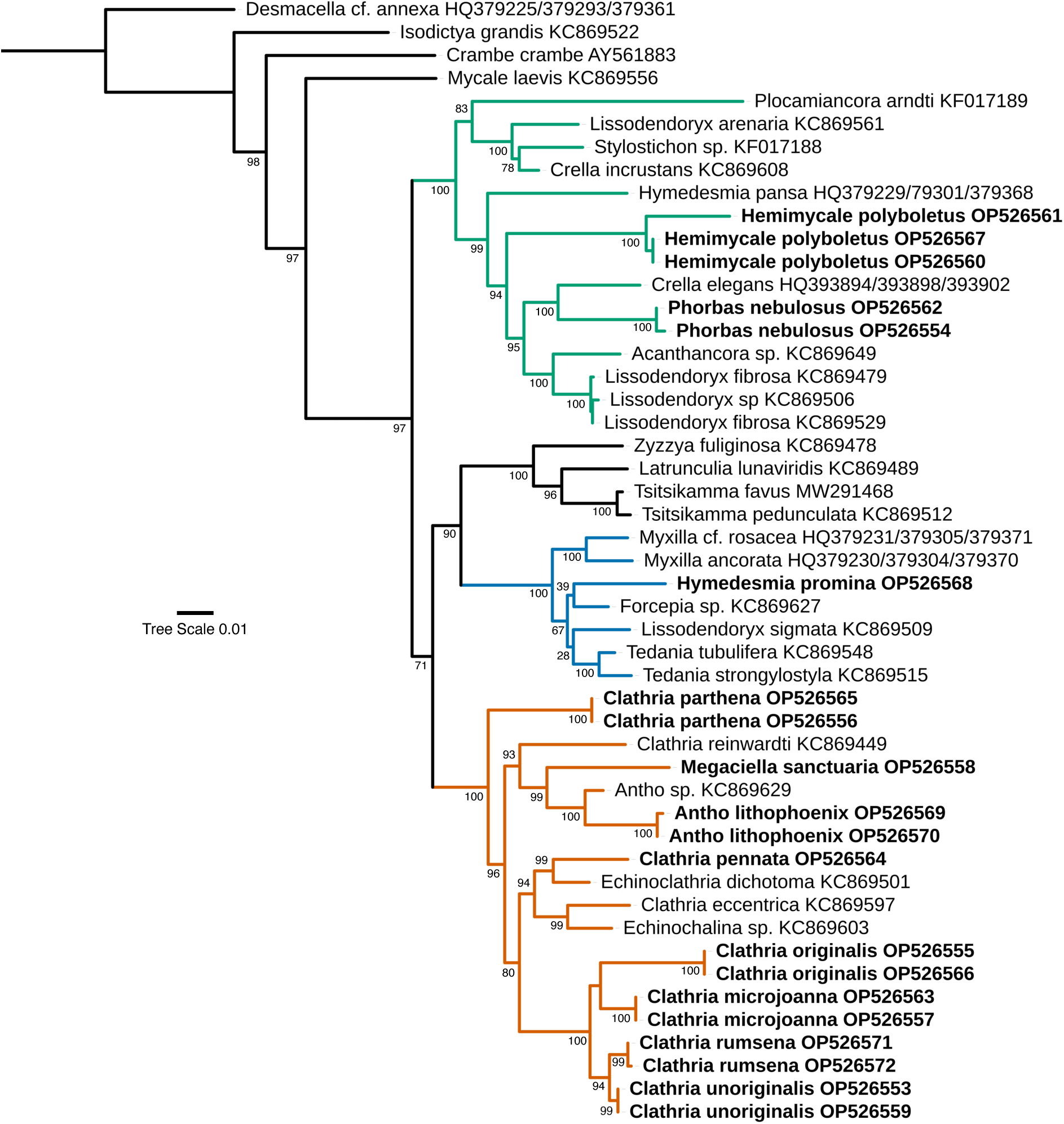
Maximum likelihood phylogeny of the *28S* locus for the Poecilosclerida. Genbank accession numbers are shown; bold indicates new sequences. Node confidence is based on bootstrapping. Scale bar indicates substitutions per site. Colors indicate clades containing new taxa, as referenced in the text.

**Figure 3.**
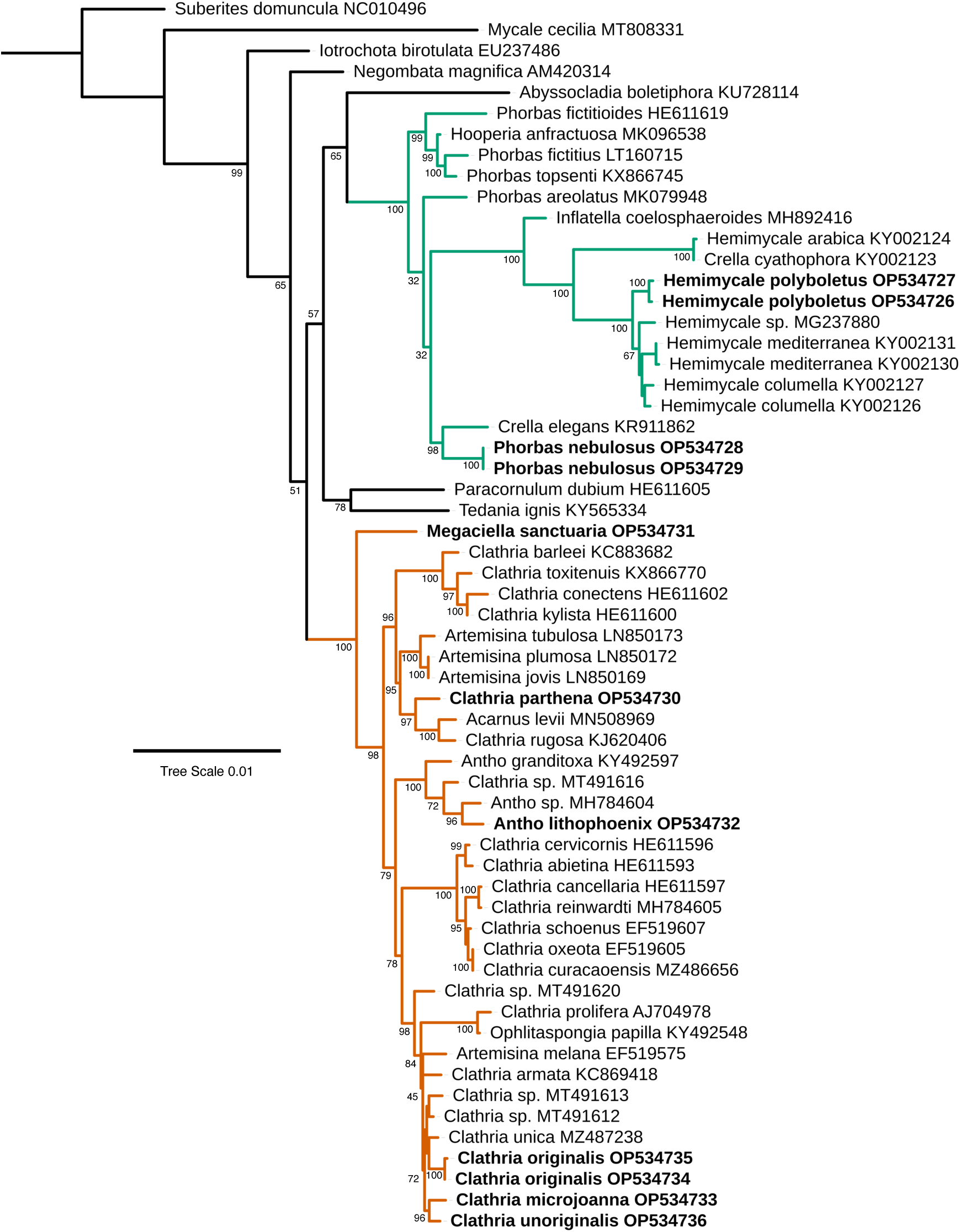
Maximum likelihood phylogeny of the *cox1* locus for the Poecilosclerida. Genbank accession numbers are shown; bold indicates new sequences. Node confidence is based on bootstrapping. Scale bar indicates substitutions per site. Colors indicate clades containing new taxa, as referenced in the text.

#### Family Hymedesmiidae

*Phorbas nebulosus sp. nov*.

Figures 2–4.

**Figure 4.**
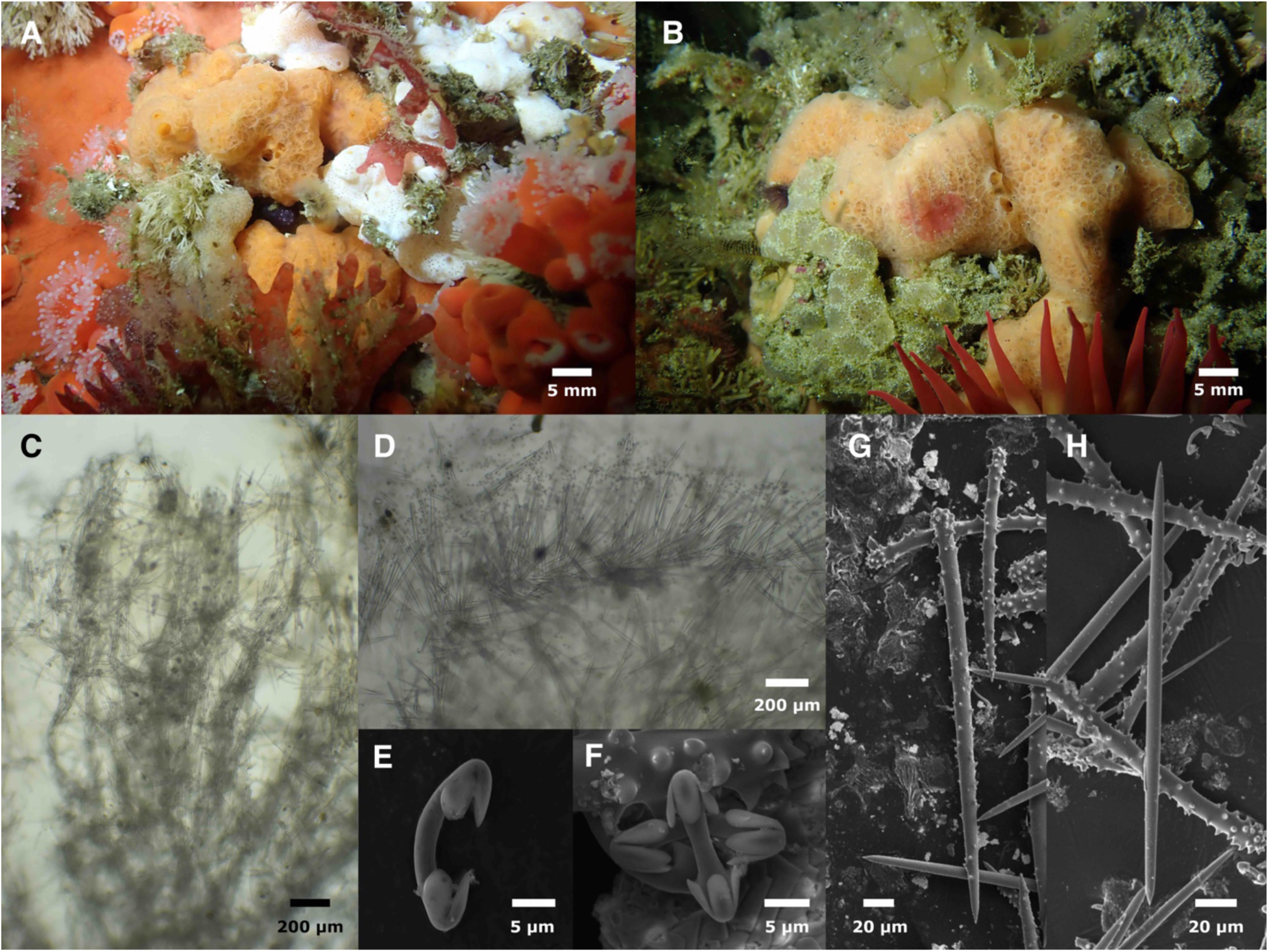
*Phorbas nebulosus*. A: Holotype in the field. B: Field photo of SBMNH700928. C: Choanosomal skeleton. D: Ectosomal skeleton. E–F: Chelae. G: Long and short acanthostyles. H: Tornote. C–H from holotype.

**Material examined.** Holotype: CASIZ236661/IZC00048467, Inner Carmel Pinnacle, (36.55910, −121.96630), 10–18 m, 8/10/21; Paratype: SBMNH700928, Wycoff Ledge, San Miguel Island, (34.02132, −120.38710), 9–19 m, 8/25/19.

**Etymology.** Named for its resemblance to supernova remnants such as the crab nebula.

**Morphology.** Thickly encrusting; holotype is 2 cm thick, paratype is 7 mm thick. Pale orange when alive, beige when preserved. Sponge surface is nearly covered in areolae, with scattered oscula also present. Areolae flush with surface, but oscula slightly raised.

**Skeleton.** Choanosomal skeleton with plumose dendritic columns of acanthostyles rising towards the surface; tracts are loose and confused, with many individual spicules also in confusion. Ectosomal skeleton of upright and tangential tornote bouquets. Chelae present throughout, but most dense in ectosome.

**Spicules.** Long acanthostyles, short acanthostyles, tornotes, arcuate isochelae. Measurements below are for both samples pooled.

Long acanthostyles: moderately spined throughout, usually most densely spined at head. 224– 283–317 x 7–11–15 μm (n=11).

Short acanthostyles: spines average larger and more dense than large acanthostyles, but with some overlap. 127–166–182 x 6–10–12 μm (n=17).

Tornotes: straight rods with both ends tapering to points; points variable, but most smoothly tapering, resembling oxeas; many appear symmetrical, while others are slightly asymmetrical, with one point more abruptly tapering than the other. 205–245–284 x 5–6–9 μm (n=40).

Arcuate isochelae: curved shaft, three alae; center ala slightly longer than side alae, and variably rounded or slightly pointed. 14–18–21 μm (n=32).

**Distribution and habitat.** Known from two samples, one from the Carmel Pinnacles and one from San Miguel Island (Southern California); both locations were natural, rocky subtidal reefs. The species is likely to be rare in the investigated region, as it is visually distinctive and has not been seen at any other site.

**Remarks.** The arcuate isochelae, areolated pore fields, plumose tracts of acanthostyles, and tangential ecotosomal brushes of tornotes lead to placement in *Phorbas*. The genetic data are unable to strongly support or refute this placement, as the genus appears highly polyphyletic (figures 2 & 3). Genetic data from the type species, *P. amaranthus* Duchassaing & Michelotti, 1864, are available at 18S, and place the type species in a poorly supported clade near *Lissodendoryx fibrosa*, *Crella elegans*, and *Hemimycale columella* (Redmond *et al*. 2013). Our phylogenies share some species in common with the published 18S phylogeny (but not *P. amaranthus*), and appear to place *P. nebulosus* sp. nov. in this same large clade, albeit closer to *Crella elegans* than to any other *Phorbas*. The genus *Crella* also has areolated pore fields, but is currently defined by having a tangential crust of spiny oxeas, anisoxeas, or styles (van Soest 2002). The genus *Crella* would therefore have to undergo substantial revision to house *P. nebulosa* sp. nov.

Two other California species are currently placed in the genus *Phorbas*: *P. californianus* (de Laubenfels, 1932) and *P. hoffmani* (Bakus, 1966). Both are easily differentiated from the new species by multiple spicular characters including the absence of chelae. The only other *Phorbas* known from the Northeast Pacific is *P. reginae* Aguilar-Camacho & Carballo 2012, from the Gulf of California, Mexico. This last species is well differentiated from the new species by multiple spicular characters including possession of tylotornotes and differently shaped chelae. *Phorbas reginae* also has much smaller acanthostyles, with the larger class averaging less than half as long as *P. nebulosus* and the small acanthostyles only a third as long as *P. nebulosus*.

Morphology also differs, with *P. reginae* being black in ethanol and lacking of visible ostia and oscules, but these differences could be due to the greater age of the *P. reginae* specimen (Aguilar-Camacho & Carballo 2012).

*Phorbas nebulosus* sp. nov. can be tentatively identified in the field based on color and being completely covered in roughly circular pore fields. *Hemimycale polyboletus* comb. nov., nom. nov. (see below) is similar, but the pore fields of that species (and *P. hoffmani)* are usually elevated on papillae. Known samples of *P. californianus* are white or blue, not orange, and the pore fields are larger and amorphous. There are also one or more undescribed species covered in circular areolae in California, similar in color to *P. nebulosus*, but these species are thinly encrusting rather than thickly encrusting and globular.

***Hemimycale polyboletus* comb. nov. nom. nov.**

Figures 2, 3, 5

**Figure 5.**
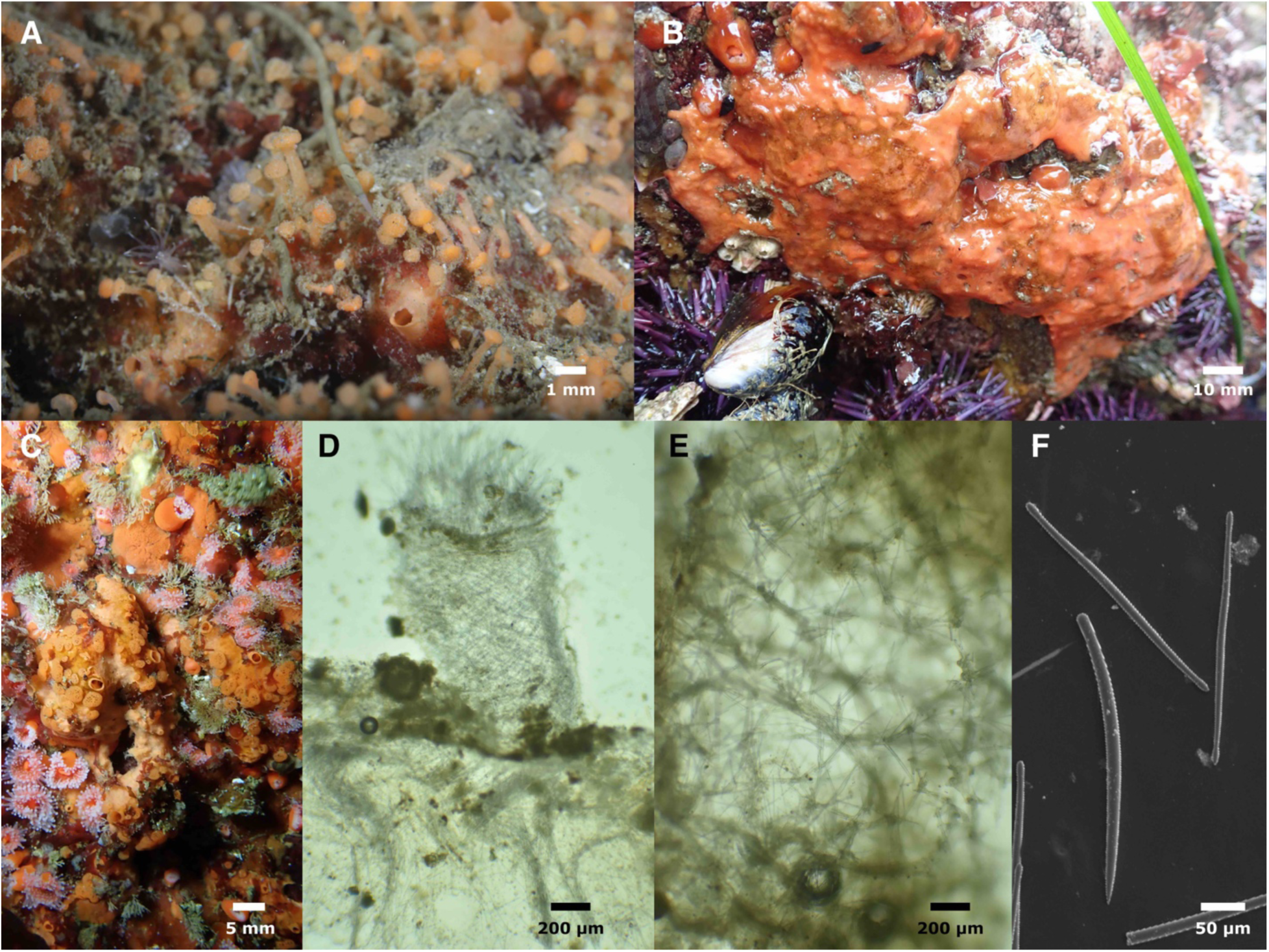
*Hemimycale polyboletus*. A: Field photo of IZC00048450. B: Field photo of SBMNH700910 exposed at low tide. C: Field photo of IZC00048449. D: Skeletal structure of choanosome and papilla, IZC00048450. E. Skeletal structure of choanosome, SBMNH700922. F: Style and subtylotes, IZC00048449.

**Synonymy.**

*Lissodendoryx (Lissodendoryx) topsenti* (Lee *et al*. 2007)

*Kirkpatrickia topsenti* (Bakus 1966)

*Tedania topsenti* (de Laubenfels 1930, 1932)

**Material examined.** IZC00048449, Inner Carmel Pinnacle, (36.55852, −121.96820), 10–24 m, 9/22/21; SBMNH700910 and SBMNH700911, Asilomar at Arena Ave, Pacific Grove, (36.62744, −121.94098), intertidal, 2/9/21; SBMNH700922, Naples Reef, Santa Barbara (34.42212, −119.95154), 6–16 m, 10/29/21; IZC00048450, Naples Reef, Santa Barbara (34.42212, −119.95154), 6–16 m, 7/15/19.

**Etymology.** Named for the many mushroom–shaped papillae decorating the surface.

**Morphology.** Encrusting, 2–30 mm thick. Apricot (light yellow–orange) or salmon (orange– pink) alive, beige when preserved. Subtidal samples have abundant areolated pore fields presented on raised, mushroom–like papillae, similar to *Cliona californiana*; scattered oscula also present. Papillae retracted in preserved samples, but still visible as circular areolae on sponge surface; papillae and oscula not visible in intertidal samples before or after preservation.

**Skeleton.** Wavy tracts of spongin cored by styles and subtylotes ascend vertically; chaotic reticulation of single spicules bound by spongin also present; reticulation is primarily styles but also some subtylotes. Ectosome with a vertical palisade of subtylotes only.

**Spicules.** Smooth styles and sinuous subtylotes/strongyles.

Styles: Of uniform thickness or with weak tyle caused by slight constriction near head end; usually curved or bent near head; pointed end sharp but often tapers in an uneven way, making the tips look roughly hewn. Less common than subtylotes/strongyles. 211–298–351 x 2–9–15 μm (n=70).

Subtylotes/strongyles: Rods with weakly swollen ends (tyles) of oblong shape; distinctly sinuous due to uneven thickness. Most with both ends swollen, but some with one end tapering to a blunt strongylote tip. 223–267–312 x 3–8–11 μm (n=90).

**Distribution and habitat.** Known from Central and Southern California (Lee *et al*. 2007), with additional unpublished reports that the sponge is present in British Columbia. Three of the samples examined here were collected near the type location (the intertidal zone at Pescadero Point, approximately 1.4 km north of the Carmel Pinnacles).

**Remarks.** Sponges in the genus *Hemimycale* have been difficult to place, as they are largely defined by traits they lack. The Systema Porifera defines the genus as “Hymedesmiidae without acanthostyles; without microscleres; megascleres strongyles and styles, not divisible into ectosomal or choanosomal spicules” (van Soest 2002). The species under discussion meets this definition. Like the type species *H. columella* (Bowerbank, 1874), it also possesses surface areolae, strengthened by upright spicules, with wavy tracts of spicules in the choanosome. The *cox1* phylogeny confirms this placement, as the species is closely related to the type species.

This species was originally assigned to *Tedania*, but it lacks onychaetes, so this was revised to *Kirkpatrickia* (Bakus 1966). *Kirkpatrickia*, however, have acanthostyles, so this assignment is not correct based on the current taxonomy. The species was then assigned to *Lissodendoryx* (Lee *et al*. 2007), but this is inconsistent with both the morphological taxonomy and the molecular phylogeny.

The name *Hemimycale topsenti* is unavailable, as another species with the same specific name, *Suberites topsenti* Burton, 1929 has already been reassigned to *Hemimycale* (Goodwin *et al*. 2019). As a result, we give this species a *nomina nova*.

*Hemimycale polyboletus* comb. nov., nom. nov. can often be identified in the field based on morphology alone. The *Cliona*-like surface morphology of this species, combined with the apricot color, are unique compared to other known species in the region. Samples exposed by the tide did not have these surface features visible, and would be much harder to identify without examination of spicules.

***Hymedesmia (Hymedesmia) promina sp. nov*.**

Figures 2 & 6

**Figure 6.**
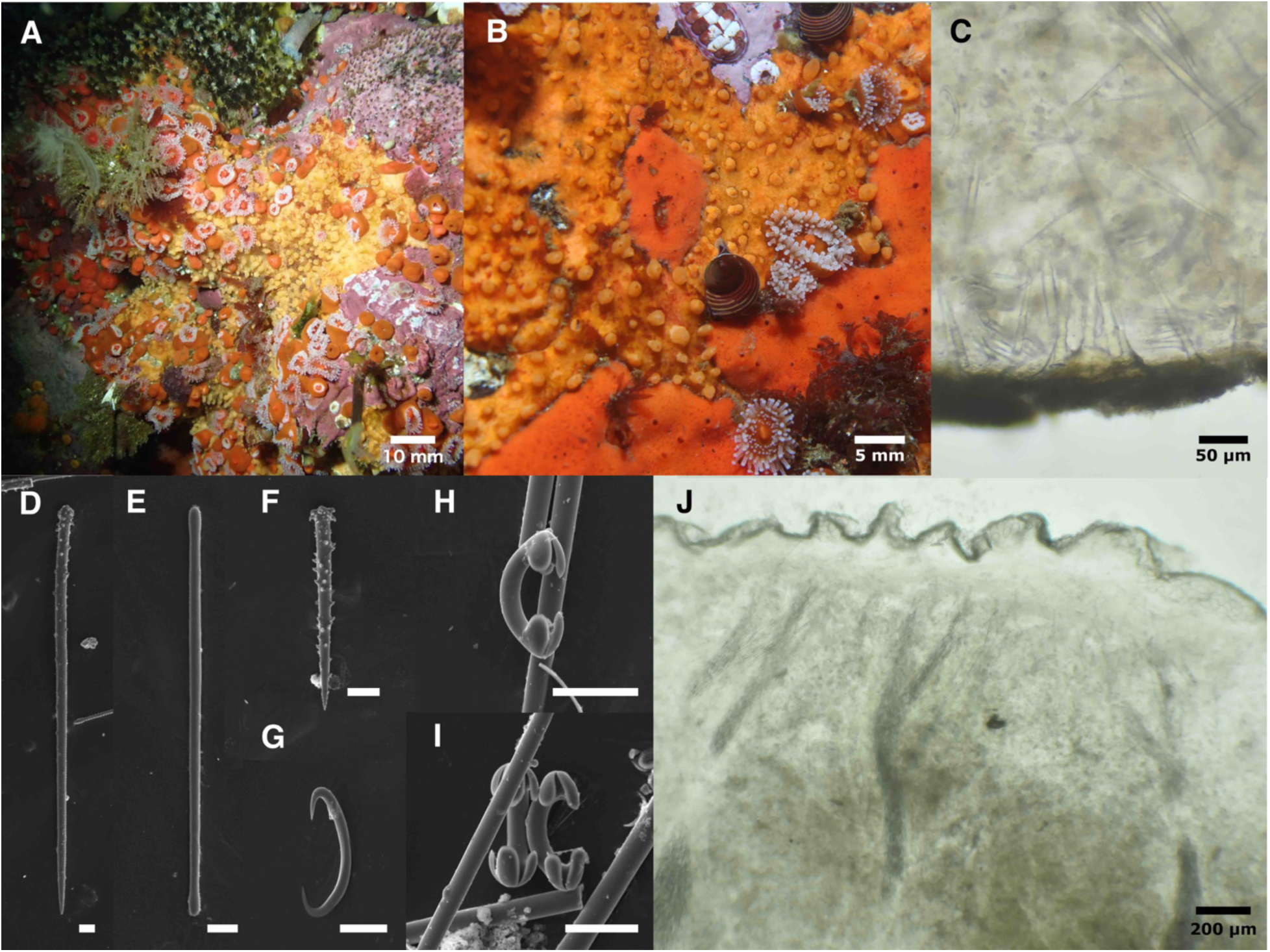
*Hymedesmia promina*. Field photos of holotype (A) and uncollected sample (B). C: Cross–section showing acanthostyles embedded in basal plate and horizontal spicules at sponge base. D: Large acanthostyle. E: Subtylote. F: Short acanthostyle. G: Sigma. H–I: Chelae. J:

Cross–section showing ascending tracts of subtylotes, with diatoms visible at sponge surface. C– J from holotype.

**Material examined.** Holotype: CASIZ236656/IZC00048451, Inner Carmel Pinnacle (36.55910, −121.96630), 10–18 m, 8/10/2021.

**Etymology.** Named for both the pinnacle on which it was found and the raised, tower-like prominences decorating the surface.

**Morphology.** Thinly encrusting, 1 mm thick, with numerous oscula atop prominences 1–2 mm in diameter; prominences occur singly, in pairs, or as ridges of several in a line. Yellowish– orange in life; turns distinctly blue in ethanol. Prominences retract when preserved but remain visible as blue mounds, with the ectosome between them appearing wrinkled and grayish. Firm and incompressible.

**Skeleton.** Basal spongin plate; some acanthostyles singly erect on substrate, with heads embedded in spongin; other acanthostyles and some subtylotes laying horizontally along substrate. Tracts of subtylotes ascend from substrate and branch, but stop before reaching ectosome; numerous microscleres scattered throughout sponge. Ectosome contains abundant diatoms and scattered chelae, but no macroscleres.

**Spicules.** Long acanthostyles, short acanthostyles, subtylotes, arcuate isochelae, sigmas.

Long acanthostyles: Densely spined in head region, sparsely spined on shaft. 316–442–513 x 11– 13–17 μm (n=17).

Short acanthostyles: Densely spined throughout; large spines on head curve towards tip, large spines on shaft curve towards head. 104–123–140 x 5–9–13 μm (n=25).

Subtylotes: Straight rods with slightly swollen, oblong tips. 244–282–404 x 3–6–12 μm (n=20).

Arcuate isochelae: Of typical shape for genus; curved shaft, three well delineated teeth, central tooth longer than lateral teeth. 27–31–37 μm (n=20).

Sigmas: C- or S-shaped, most rotated such that they are not in a single plane. 41–53–65 μm (n=23).

**Distribution and habitat.** Known only from the Carmel Pinnacles. Numerous individuals seen on initial dive, but only one sample taken; no individuals seen on second dive.

**Remarks.** *Hymedesmia* are thinly encrusting Hymedesmiidae with acanthostyles singly erect on the substrate and bundles of tornotes in the choamosome and/or ectosome (van Soest 2002). Genetic data indicate the genus is polyphyletic (Redmond *et al*. 2013); the ‘hymedesmoid’ skeletal morphology is known to have evolved multiple times outside the family, so it is not surprising that it has also evolved repeatedly within the family (van Soest 2002). No genetic data is yet available for the type species of *Hymedesmia*, and data from few species are available at *28S* or *cox1*. Surprisingly, however, *H. promina* appears to be more closely related to species placed in the Coelosphaeridae and Myxillidae than to other Hymedesmiidae (figure 2). As this species is clearly in *Hymedesmia* based on the morphological taxonomy, reconciling the taxonomy with the evolutionary history of hymedesmoid sponges must await genetic data from more species and a systematic revision.

*Hymedesmia promina* is easily differentiated from other described species known from the region. *Hymedesmia (Stylopus) arndti* (de Laubenfels, 1930) is the only *Hymedesmia* described from the northeast Pacific, though there are also unpublished reports of *H. (Stylopus) primitiva* Lundbeck, 1910 occurring in British Columbia; neither species has chelae. An additional 11 sponges have been referred to as “*Hymedesmia* sp.” in various California surveys and remain undescribed (Lee *et al*. 2007). Of these, an intertidal sample from the Farallon Islands in Northern California (Klontz 1989) is the only potential match to *H. promina* sp. nov.. These previous undescribed sponges are vouchered but were not examined as part of this work.

Further work is required before this species can be confidently identified based on gross morphology in the field. Thinly encrusting orange sponges are numerous in California, but it is possible that the tower-shaped oscula of *H. promina* sp. nov. are diagnostic.

Family Microcionidae

***Clathria (Microciona) unoriginalis sp. nov*.**

Figures 2, 3, 7

**Figure 7.**
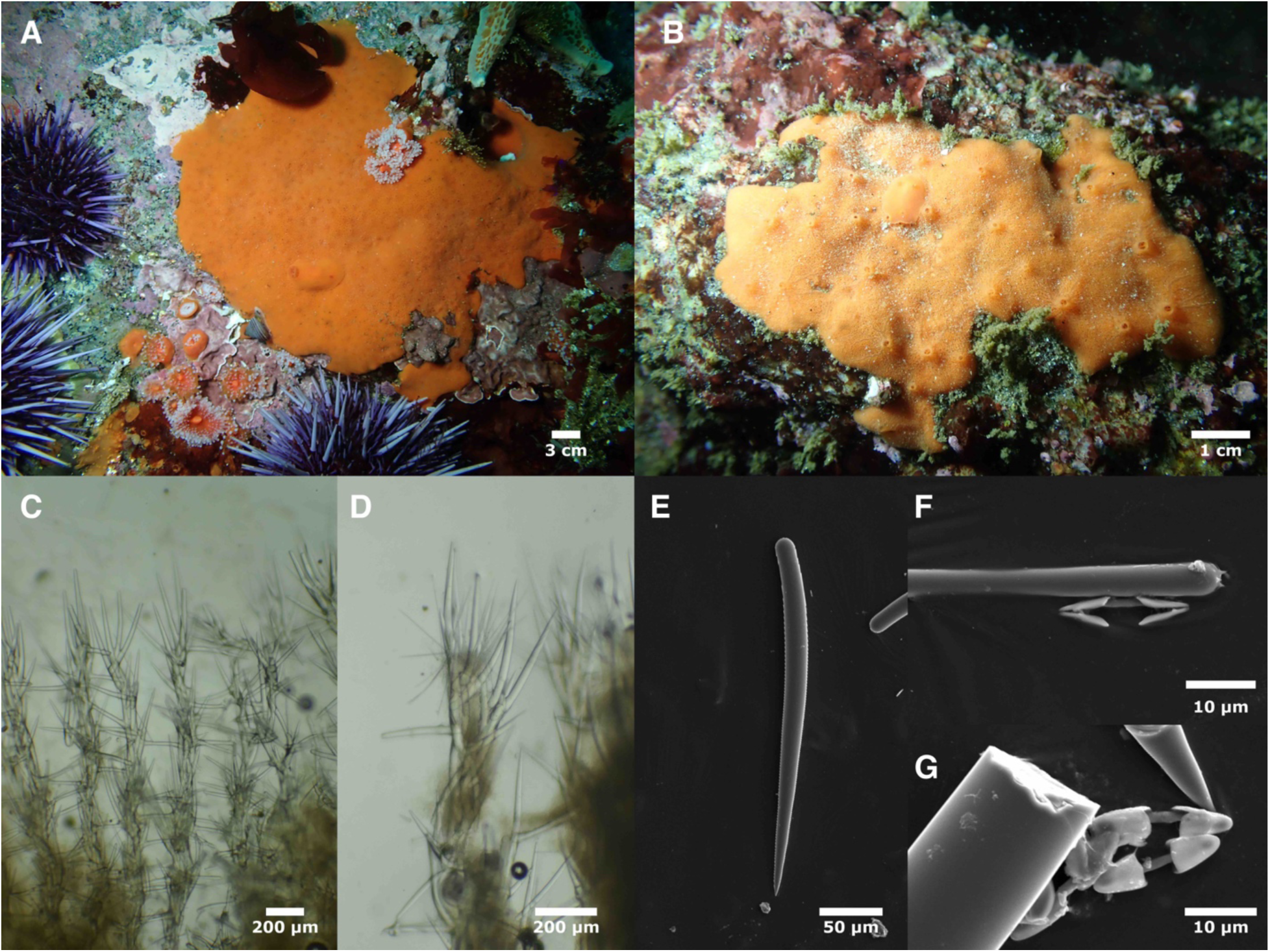
*Clathria unoriginalis*. Field photos of holotype (A) and IZC00048439 (B). Cross sections showing choanosomal skeleton (C) and ectosomal skeleton (D). E: Thick style. F: Head of thin style and side view of chela. G: Chelae. C–G from holotype.

**Material examined.** Holotype: CASIZ236653/IZC00048440, Inner Carmel Pinnacle, (36.55852,-121.96820), 10–24 m, 9/22/21; paratypes: SBMNH700906, Arroyo Quemado Reef, Santa Barbara, (34.46775, −120.11905), 7–11 m, 1/7/20; IZC00048439, Cannery Row, Monterey, (36.61798, −121.89780), 9–16 m, 9/21/21; IZC00048438, 1000 Steps Reef, Santa Barbara, (34.39472, −119.71347), 4–6 m, 7/12/19; SBMNH700912, Carmel Point, Carmel, (36.54488, −121.93300), intertidal, 2/7/21.

**Etymology.** Named for its similarity to the sympatric congener, *Clathria originalis* (de Laubenfels, 1930).

**Morphology.** Thinly encrusting, 1–2 mm thick; bright orange to orange-red in life, fades to beige in ethanol. Surface covered in minute pores; numerous, evenly-spaced round oscula, each 1–2 mm across, present in larger samples; oscula with slightly raised rims, usually surrounded by a vein-like network of subsurface channels.

**Skeleton.** Columns of spongin cored with large styles; coring styles close to vertical, leading to fairly compact columns, but some styles with tips free and angled out, making columns slightly plumose. Coring spicules at the top of columns flare out into bouquets. Columns chaotically bridged by single, horizontal spicules in spongin, making skeleton ladder-like. Thin styles present as disorganized clumps within bouquets of large spicules at the top of columns. Few chelae seen in sections; all were near ectosome. No toxas seen in sections.

**Spicules.** Thick styles, thin ectosomal subtylostyles, palmate isochelae, toxas.

Thick styles: gently curved and gradually tapering. Thin (potentially immature) styles in this category are differentiated from thin subtylostyles by the absence of a tyle, more pronounced curvature, and more gradual taper. In one sample (IZC00048438), thick styles were mostly subtylote and some bore minute dimples on head; other samples had few or no styles that were subtylote and no dimpling on heads. Holotype 203–328–525 x 16–22–29 μm (n=53); all samples pooled 135–253–525 x 7–17–29 μm (n=197); lengths quite variable across samples, but variation not partitioned by geographic region; mean values per sponge from 207 to 328 μm.

Thin ectosomal subtylostyles: many subtylote, usually with small spines on heads. Straight or slightly curved. In some samples the thin styles averaged shorter than thick styles, but in other samples this was reversed. Holotype: 163–268–452 x 3–5–6 μm (n=21); all samples pooled 133– 291–506 x 2–4–6 μm (n=82); mean lengths per sponge from 210 to 325 μm.

Palmate isochelae: holotype 14–19–21 μm (n=15); all samples pooled 13–17–21 μm (n=44).

Toxas: none seen in holotype; common in one sample (IZC00048438); rare in other samples. All samples pooled: 44–113–206 μm (n=26).

**Distribution and habitat.** Of the five known samples, 2 were collected on the Santa Barbara coast (Southern California) and three are from around the Monterey Peninsula (Central California). Intertidal to at least 10 m depth on natural reefs where kelp forest is found.

**Remarks.** The genus *Clathria* is well represented in the Northeast Pacific, with 11 previously described species in the region. Only two of these species lack acanthostyles: *C. originalis* (de Laubenfels, 1930) and *C. pennata* (Lambe, 1895); of these, only *C. originalis* has chelae. *Clathria unoriginalis* is very similar to *C. originalis* in terms of morphology, skeletal structure, and spiculation, but these species can be distinguished by the longer styles of *C. unoriginalis.* The styles of *C. originalis* were originally described as 150–155 μm in length; a more recent publication lists the thick styles as 130–165 μm and the thin styles as 90–200 μm (Lee *et al*. 2007). Preliminary analysis of 13 *C. originalis* samples collected by the authors finds that the mean values for each sponge vary from 102–155 μm for thick styles and 100–127 μm for thin styles. Though the values for *C. unoriginalis* are highly variable, all samples average longer than *C. originalis*: 207–328 μm for thick styles and 210–325 μm for thin styles. These long styles are more similar to *C. pennata*, which opens the possibility that the presence of chelae is variable in *C. pennata.* This is refuted by the DNA data, which support species status for all three species (figures 2 & 3). The encrusting morphology and skeletal structure of this species are consistent with assignment to the subgenus *Microciona*.

The World Porifera Database currently lists an additional *Clathria* species for the region that lacks acanthostyles: *C. hartmani* (Simpson, 1966). This name is a junior synonym of *C. originalis*. When de Laubenfels first published descriptions of the encrusting orange sponges of the Monterey Peninsula, he mentioned that some *C. pennata* have chelae, and said these forms may be a different variety (de Laubenfels 1927). After additional study, de Laubenfels named the forms with chelae *C. originalis* (de Laubenfels 1930, 1932). Simpson (1966) later published the name *C. hartmani* for the type with chelae, citing the 1927 paper, and apparently not realizing that the name *C. originalis* had already been applied. As a result, all later authors working in the region correctly considered *C. hartmani* a junior synonym of *C. originalis* (Bakus & Green 1987; Hartman 1975; Lee *et al*. 2007).

It was noted by de Laubenfels that *C. pennata* occurs closer to the high tide mark than any other California sponge, with *C. originalis* tending to occur lower in the intertidal and also present in the shallow subtidal (de Laubenfels 1932). Our recent collections have found all three species in both the subtidal and intertidal, but with different frequencies: 4 of 5 *C. unoriginalis* sp. nov. were found in the subtidal, but only 2 of 13 *C. originalis* and 1 of 8 *C. pennata* were found subtidally. Subtidal samples of the latter two species were found only on shore dives in very shallow water. The precise elevation/depth of each sample was not recorded, but we hypothesize that *C. unoriginalis* sp. nov. averages depths below the other two species — though they clearly overlap.

*Clathria unoriginalis* sp. nov. cannot be identified in the field. In addition to the sympatric species of *Clathria*, several species of thinly encrusting *Antho* occur in the same habitats. Average differences in color and texture among species have been noted in specific locations, but overall, variability has thwarted all attempts to find useful field marks.

***Clathria (Microciona)* rumsena sp. nov.**

Figures 2, 8

**Figure 8.**
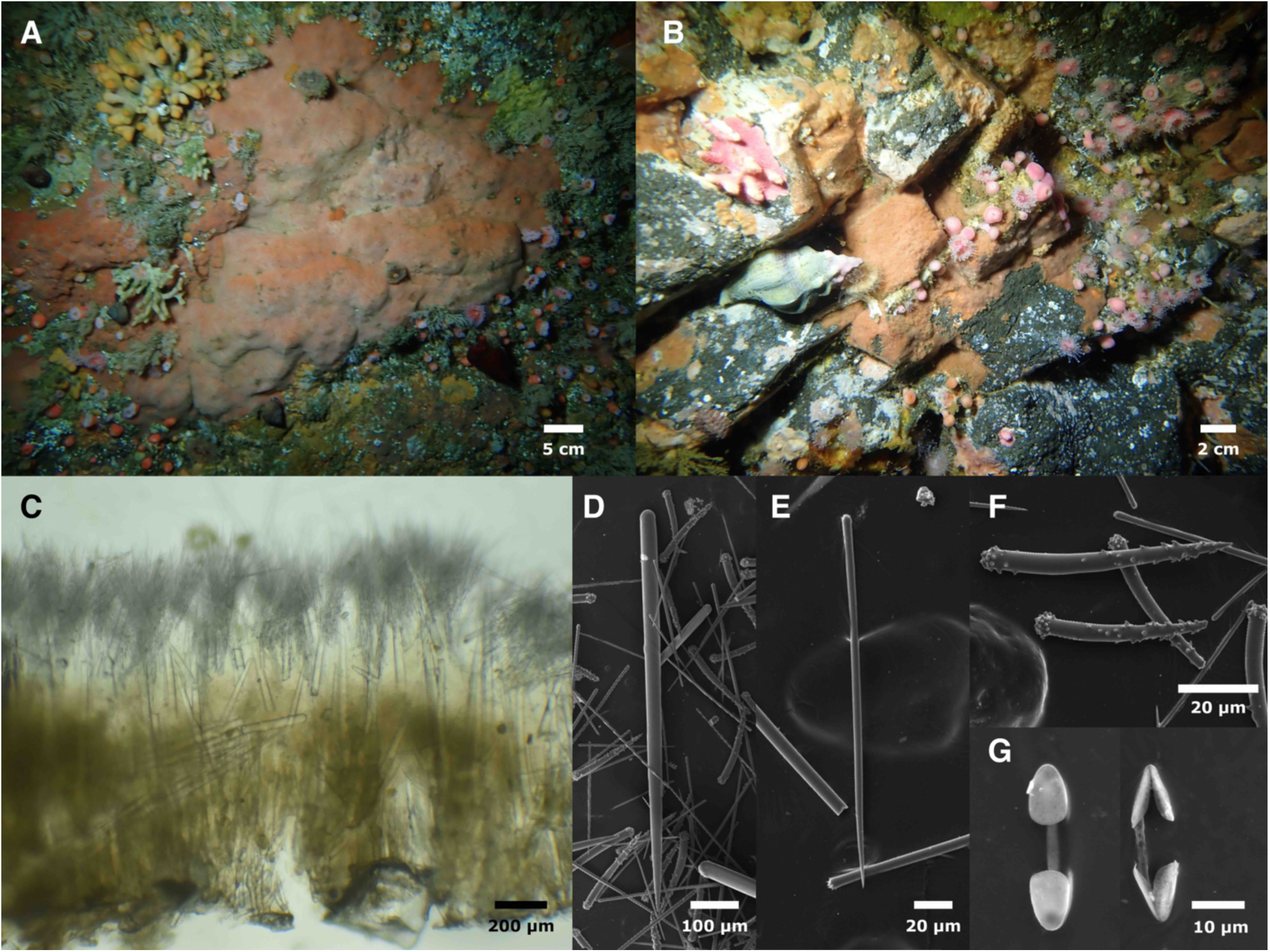
*Clathria rumsena*. Field photo of holotype (A) and IZC00048437 (B). C: Cross section. D: Thick style. E: Thin style. F: Acanthostyles. G: Chelae. C–G from holotype.

**Material examined.** Holotype: CASIZ236652, Inner Carmel Pinnacle, (36.55852, −121.96820), 10–24 m, 9/22/21; paratype: IZC00048437, Fire Rock, Pescadero Point, Carmel, (36.55898, − 121.95110), 10–22 m, 8/10/22.

**Etymology.** Named in honor of the Costanoan Rumsen Carmel Tribe, whose ancestral lands include those surrounding Monterey and Carmel bays.

**Morphology.** Thinly encrusting, 1–2 mm thick; light orange to peach colored alive; fades to beige in ethanol. Surface of holotype bearing evenly-spaced round oscula surrounded by a network of transparent surface channels.

**Skeleton.** Thick styles with heads embedded in basal spongin plate and nodes of spongin arising from it. Acanthostyles echinating spongin nodes. Thin styles present as dense, upright bouquets at sponge surface.

**Spicules.** Thick styles, thin straight styles, acanthostyles, palmate isochelae.

Thick styles: curved and tapering; some are weakly subtylote and have weakly spined heads. Holotype had a few capsule or strongyle-shaped spicules that appeared to be aberrant styles with two rounded ends. Holotype 384–744–1095 x 17–29–38 μm (n=35), both samples combined 384–729–1095 x 7–28–38 μm (n=40)

Thin straight styles: straight; some have very weak tyles, but many unadorned. Holotype 158– 292–637 x 3–5–10 μm (n=138), both samples combined 158–287–637 x 3–5–10 μm (n=155)

Acanthostyles: heads densely covered in round bump–like spines; spines on shaft more pointed, curving towards head, most dense near tips. Holotype 121–188–287 x 11–15–20 μm (n=30), both samples combined 112–188–381 x 8–15–27 μm (n=48).

Palmate isochelae: holotype 12–15–17 μm (n=21), both samples combined 12–16–20 μm (n=52).

**Distribution and habitat.** Known from two samples, both collected on natural rocky reefs offshore of Pescadero Point, Carmel Bay, California.

**Remarks.** *Clathria rumsena* is differentiated from all other *Clathria* in the region by having longer styles of all types (thick styles, thin styles, and acanthostyles). The most similar species is *Clathria asodes* (de Laubenfels, 1930), a thinly encrusting yellow species. In addition to the color difference, the maximum lengths of all *C. asodes* spicule types — including chelae — are less than the average lengths for *C. rumsena* (de Laubenfels 1932; Lee *et al*. 2007). All other thinly encrusting *Clathria* in the region have qualitative as well as quantitative differences, such as lacking chelae (*C. pseudonapya* (de Laubenfels, 1930)) or having arcuate chelae (*C. brepha* (de Laubenfels, 1930), and *C. spongigartina* (de Laubenfels, 1930)). Thickly encrusting *Clathria* from the region differ in additional ways as well, such as lacking spines on large styles; these species are also genetically differentiated as shown in the phylogenies (figures 2 & 3).

The encrusting morphology and skeletal structure of this species are consistent with assignment to the subgenus *Microciona*. The subgenus *Thalysias* was also considered: this subgenus is defined by having two size classes of thin styles, with larger forms subectosomal, supporting brushes of small styles in the ectosome (Hooper 2002b). The thin styles of *C. rumsena* are found in a large size range, but have a continuous distribution with a single mode, and no evidence of a multi-layered ectosomal skeleton could be found.

*Clathria rumsena* sp. nov. cannot be identified in the field. In addition to the sympatric species of *Clathria*, several species of thinly encrusting *Antho* occur in the same habitats. Some average differences in color, patterning, and texture have been noted in specific locations, but overall, variability frustrates attempts to identify thinly encrusting red/orange sponges in the field in California.

***Clathria (Microciona) microjoanna* (de Laubenfels, 1930)**

Figures 2, 3, 9

**Figure 9.**
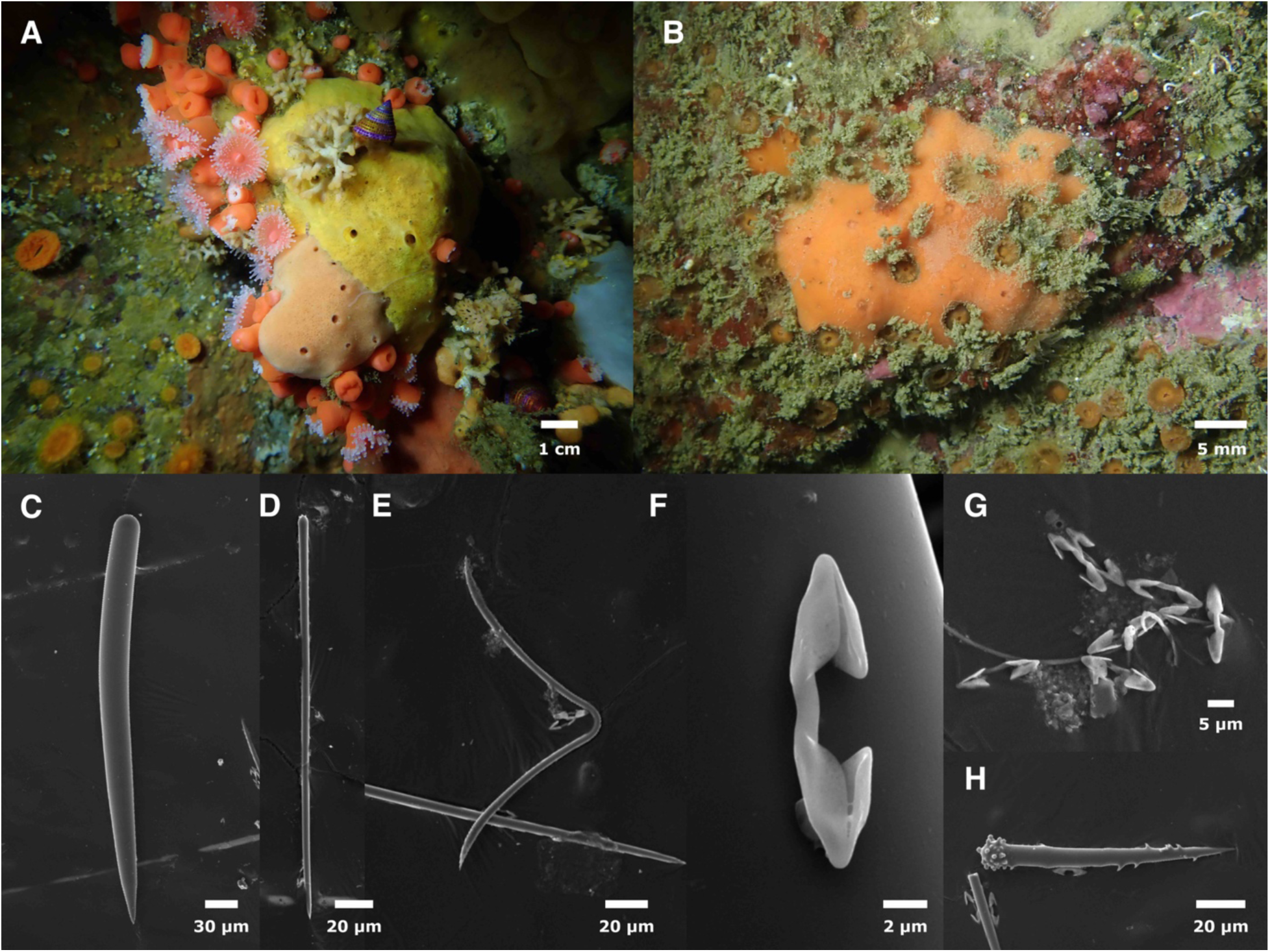
*Clathria microjoanna*. A: Field photo of IZC00048435 (peach, bottom); yellow sponge is *Antho lithophoenix* sample IZC00048434. B: Field photo of TLT477. C: Thick style. D: Thin subtylostyle. E: Toxa. F–G: Chelae. H: Acanthostyle. C–H from IZC00048435.

**Synonymy.**

*Microciona microjoanna* (de Laubenfels 1930, 1932)

**Material examined.** IZC00048435, Inner Carmel Pinnacle, (36.55852,-121.96820), 10–24 m, 9/22/21; NHMLA16556, Hawthorne Reef, Los Angeles, (33.74714, −118.42090), 18–24 m, 8/23/19; TLT477, Otter Cove, Pacific Grove, (36.62920, −121.92031), 7–12 m, 11/24/19.

**Morphology.** Thickly encrusting, firm, with prominent oscules. Samples investigated here were vivid orange to pale orange-pink alive; beige when preserved. The species has also been reported to occur in scarlet (Lee *et al*. 2007). Two of the three specimens investigated here were found in an apparently quiescent state, with oscules closed by a membrane.

**Skeleton.** Not investigated; previously reported as having plumose tracts sparsely echinated by acanthostyles, with thin styles grouped at surface (Lee *et al*. 2007).

**Spicules.** Thick styles, thin subtylostyles, acanthostyles, palmate isochelae, toxa.

Thick styles: smooth, gently curved, and tapering gradually to sharp points; thin (potentially immature) styles in this category are differentiated from thin subtylostyles based on more pronounced curve and more gradual taper. 241–314–408 x 7–17–25 μm (n=57).

Thin subtylostyles: straight or very slightly curved; sometimes with a very slight swelling at head, but this is often missing. Heads microspined. 139–213–349 x 2–5–7 μm (n=43).

Acanthostyles: uncommon. Heads densely spined, shafts sparsely spined; spines fairly large and variously knob-like or curved; spines on shaft curve towards head. 95–103–111 x 7–8–9 μm (n=12).

Palmate isochelae: 9–14–22 μm (n=79).

Toxa: with weakly microspined tips. 41–176–385 μm (n=12).

**Distribution and habitat.** Known from Southern California to Washington, from the intertidal to deep water (Bakus 1966; Lee *et al*. 2007). Type location is the intertidal zone at Pescadero Point, just onshore of the Carmel Pinnacles sample investigated here. Recent collections from diving depths indicate that this species is probably more common around the Monterey Peninsula than in Southern California.

**Remarks.** The spicules of this sponge are quite similar to the sympatric *Clathria (Microciona) parthena* (de Laubenfels, 1930). These species have quantitative differences in the sizes of the chelae, styles, and toxas, as reported in the original description and below. Lee *et al*. (2007) report that the toxas are diagnostic, with toxas in *C. parthena* not exceeding 100 μm; we have found toxas over 300 μm in *C. parthena*, though the average size is over twice as long in *C. microjoanna* (average shape also differs, see below). The chelae make an easier diagnostic character, with *C. microjoanna* chelae lengths averaging 12–16 μm per sponge, while *C. parthena* chelae average 23–26 μm per sponge. *Clathria parthena* also look different when alive, as described below.

*Clathria microjoanna* are unlikely to be identifiable in the field. With few live *C. microjoanna* photographed to date, potential field marks remain to be determined, but spicule or genetic identification will likely remain necessary even with additional data. Preliminary indications are that this species may often be thicker than many sympatric orange sponges (up to 2 cm thick), with subtle surface patterning (compared to *Dragmacidon mexicanum*), a fairly uniform surface (flat with regularly spaced and consistently-sized oscula compared to *Antho lithophoenix*), and sometimes a pinkish tinge (not yet known in other local species). These characteristics might help target collections of this species for later spicule confirmation.

***Clathria (Microciona) parthena* (de Laubenfels, 1930)**

Figures 2, 3, 10

**Figure 10.**
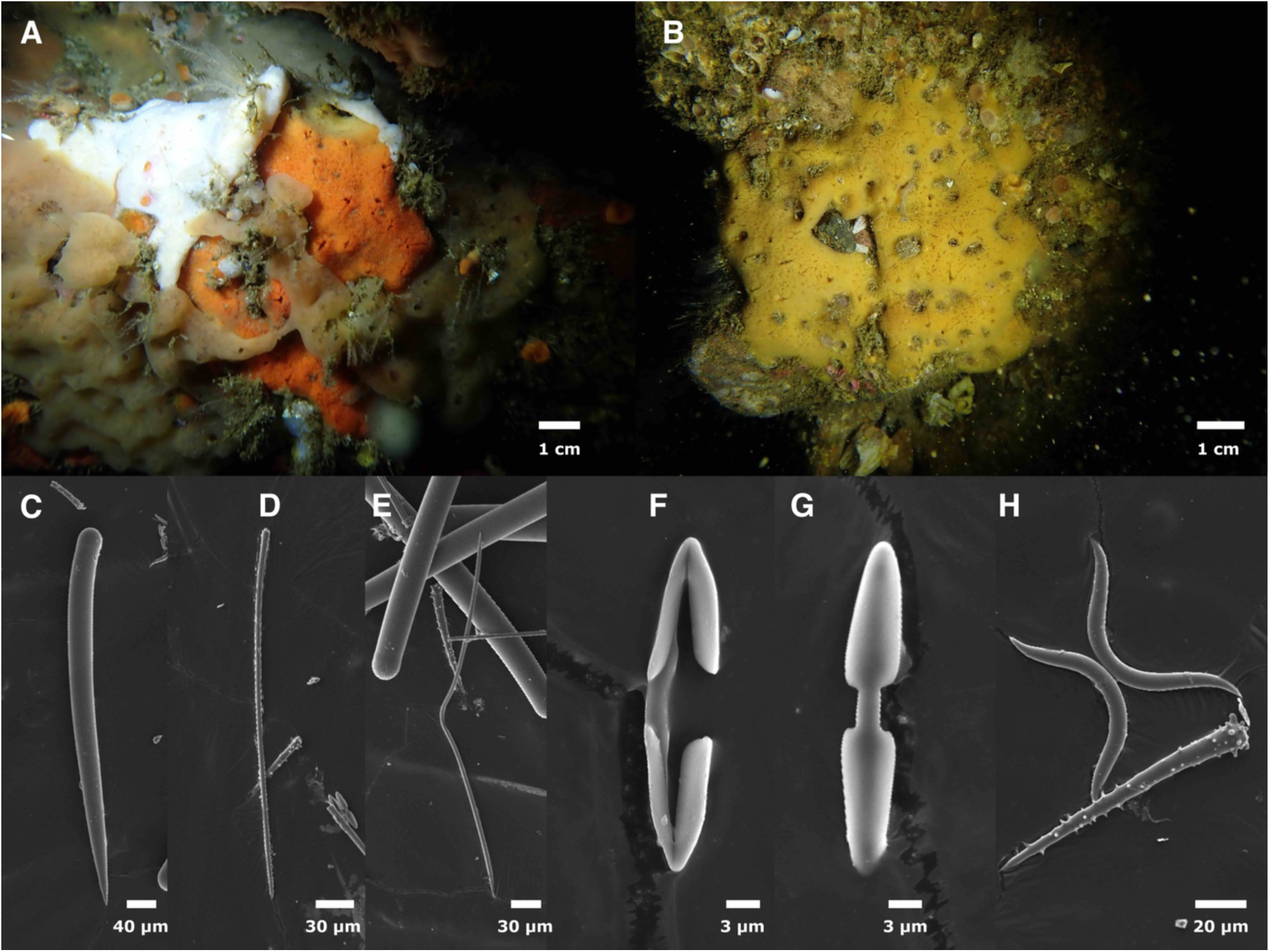
*Clathria parthena*. Field photos of IZC00048436 (A) and TLT543 (B). C: Thick style. D: Thin subtylostyle. E: Large toxa with low arch; acanthostyle and head spines on thick styles also visible. F–G: Chelae. H: Toxa with high arch and acanthostyle. C–H from IZC00048436.

**Synonymy.**

*Microciona parthena* (de Laubenfels 1930, 1932)

**Material examined.** IZC00048436, Inner Carmel Pinnacle (36.55852, −121.96820), 10–24 m, 9/22/21; TLT543, The Suburbs, Santa Cruz Island, (34.05291, −119.58309), 6–13 m, 1/19/20; TLT223, Arroyo Quemado Reef, (34.46775, −120.11905), 7–11 m, 7/29/19.

**Morphology.** Most samples are vivid orange but one mustard yellow; all fade to beige in ethanol. Columns of styles are prominent in live sponges; these combine with irregular shape and thickness to give sponges a distinctly roughened appearance. Oscula prominent, approximately 2 mm in diameter, often atop low mounds; inconsistently spaced; often irregular in shape (some round and some oval), with vein-like channels converging upon them.

**Skeleton.** Not investigated; previously reported as having plumose tracts sparsely echinated by acanthostyles, with thin styles grouped at surface (Lee *et al*. 2007).

**Spicules.** Thick styles, thin subtylostyles, acanthostyles, palmate isocheleae, toxa.

Thick styles: unspined, gently curved, and tapering gradually to sharp points; some with minute head spines; thin (potentially immature) styles in this category are differentiated from thin subtylostyles based on more pronounced curve and more gradual taper. One capsule-shaped spicule was interpreted as an aberrant style. 77-120–404 x 8–21–39 μm (n=77).

Thin subtylostyles: straight or slightly curved, some have weakly swollen heads; microspined on head. 204–321–476 x 2–4–7 μm (n=43).

Acanthostyles: well spined on head and shaft, head spines curve toward shaft and shaft spines curve towards head. 65–74–100–124 x 4–7–10 μm (n=65).

Palmate isochelae: Distinctly long and thin compared to other California Microcionidae; 18–26– 33 μm (n=57).

Toxa: Majority are distinctly thick with a high arch, resembling a handlebar mustache (figure XH); a minority are thinner, and some of these are large, flat, and have a low arch (figure XE); 11–60–367 μm (n=107).

**Distribution and habitat.** Previously reported as occurring throughout California, from the intertidal to 45 m (Lee *et al*. 2007). Recent surveys of diving depths in Southern and Central California found this species at 17% of subtidal reefs, evenly dispersed throughout the region. Often numerous when present at a site.

**Remarks.** As discussed above in the *Clathria microjoanna* section, the spicules of this species are qualitatively similar to that species, but easily differentiated quantitatively. In addition to the sizes of spicules, these species are differentiated by the shape of the chelae (elongated in *C. parthena*) and the shape of the toxas (most with a handlebar mustache appearance in *C. parthena*).

This sponge is often identifiable in the field due to its distinctly roughened, uneven appearance. Without spicule or DNA confirmation, however, it is likely that some sponges of this species would be confused with other orange sponges such as *Antho lithophoenix* or *Clathria microjoanna*.

***Antho (Jia) lithophoenix* (de Laubenfels, 1927)**

Figures 2, 3, 11

**Figure 11.**
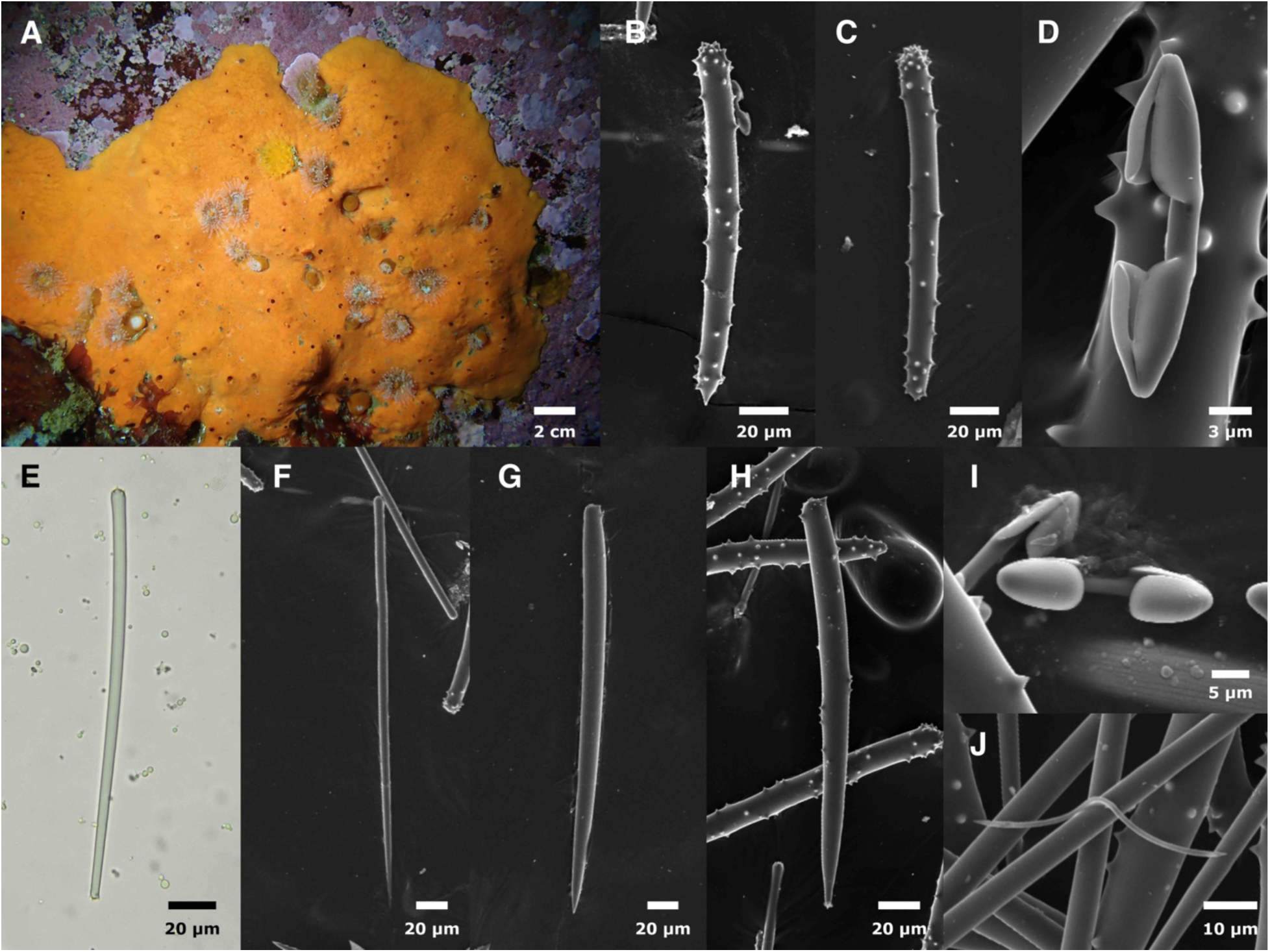
*Antho lithophoenix.* A: field photo of IZC00048433 (see also figure 9A for field photo of IZC00048434). B: choanosomal acanthostyle. C: choanosomal acanthostrongyle. D, I: Chelae. E: ectosomal strongyle. F: ectosomal style. G–H: subectosomal acanthostyles. J: toxa. B–J from IZC00048433.

**Synonymy.**

*Antho (Antho) lithophoneix* (Lee *et al*. 2007)

*Isociona lithophoenix* (de Laubenfels 1932)

*Plocamia lithophoenix* (de Laubenfels 1927)

**Material examined.** IZC00048433, Inner Carmel Pinnacle (36.55910, −121.96630), 10–18 m, 8/10/2021; IZC00048434, Inner Carmel Pinnacle, (36.55852,-121.96820), 10–24 m, 9/22/21; TLT28, Coal Oil Point, Santa Barbara (34.40450, −119.87890), 3–8 m, 4/24/19.

**Morphology.** Samples from the Carmel Pinnacles were thickly encrusting, 1–3 cm thick; comparative material from Coal Oil Point thinly encrusting, 3–5 mm thick. Samples examined here were vivid orange or yellow in life, beige in ethanol; samples previously described as brick red or scarlet (Lee *et al*. 2007). Thickness uneven, surface rugose. Oscules prominent and unevenly distributed on sponge surface; concentrated especially on high points.

**Skeleton.** Reticulation of stubby acanthostyles bound by considerable spongin at nodes. Near the sponge surface, larger styles are attached by their heads to the reticulation, singly or in plumose brushes, tips pointing up. Thin ectosomal styles and strongyles in brushes at surface. Chelae concentrated at surface but seen throughout.

**Spicules.** Choanosomal acanthostyles (with some acanthostrongyles), large subectosomal acanthostyles, thin ectosomal styles, thin ectosomal strongyles, palmate isochelae, toxas. Crocae previously reported as rare in holotype but not seen in samples examined (van Soest *et al*. 2016).

Choanosomal acanthostyles: spiny over entire surface, but most densely at head. Most were clearly styles, with a rounded head end, thick, non-tapering shafts, then an abrupt, sharp point at the other end. A minority are strongyles, with both ends rounded; these were more common in the Coal Oil Point sample than in the Pinnacles samples, where they are nearly absent. Carmel samples 131–152–166 x 7–11–14 μm (n=87); spicules from Southern California sample 15% smaller, 116–131–149 x 4–7–11 μm (n=34).

Large subectosomal acanthostyles: sparsely spined, most densely spined at head; some completely unspined. Longer and more tapering than choanosomal styles, but difficult to separate. Carmel samples 172–224–346 x 5–13–20 μm (n=33).

Thin ectosomal styles: sometimes slightly subtylote but often straight; heads with minute spines; tapering to sharp points. Carmel samples 162–259–328 x 3–5–7 μm (n=31).

Thin ectosomal strongyles: similar to thick ecotomosal styles, but lack a sharp point and have minute spines at both ends. Many appear to taper slightly, and therefore resemble modified styles. Roughly equal in abundance to thin ectosomal styles. Carmel samples 134–160–206 x 3– 5–6 μm (n=33).

Palmate isochelae: Carmel samples 17–21–26 μm (n=48).

Toxas: Carmel samples 38–94–212 μm (n=18).

**Distribution and habitat.** Lee *et al*. (2007) lists the distribution of this species as Central and Southern California, intertidal to 39 m. Consistent with that range, recent collections by the authors have found this species from the intertidal and the subtidal throughout Central and Southern California. However, due to spicule variability and potential confusion with *Antho illgi* (Bakus, 1966), most of these samples were not analyzed here, and will be included in a comprehensive future review (see Remarks below). The Carmel Pinnacles sponges are from near the type location: the intertidal zone in Pacific Grove.

**Remarks.** This species was first described as having acanthotylotes that approached acanthostrongyles (de Laubenfels 1927); de Laubenfels later modified this to acanthostyles and acanthostrongyles (de Laubenfels 1932); reanalysis of the holotype found that the vast majority were acanthostyles, and only occasionally were they found as acanthostrongyles (van Soest *et al*. 2016). This reanalysis of the holotype also found that a substantial minority of the thin ectosomal spicules were strongylote, not stylote, and found rare J-crocae. The freshly collected samples investigated here are a very good match for the reanalysis of the holotype, qualitatively and quantitatively. The only exception are the J-crocae, which were not found. We therefore feel confident assigning these samples to *Antho lithophoenix*.

We have collected additional samples from throughout Southern and Central California that are also likely this species, but we do not include them here because a thorough review is needed. Preliminary results indicate that some have almost entirely acanthostyles (like the holotype and samples from the Carmel Pinnacles), others are a mix of acanthostyles and acanthstrongyles, and some are almost entirely acanthostrongyles. Samples with only acanthostrongyles still have large, sparsely-spined acanthostyles in the subectosomal region. These samples could be considered *Antho illgi* (Bakus, 1966), as this species was differentiated from *A. lithophoenix* based on the acanthostrongyles and acanthostyles being in distinct size ranges. *Antho illgi* was described from the San Juan Islands, Washington; it was later reported from California, but with data that seem more consistent with *A. lithophoenix*. A comprehensive review of these samples will be presented in a later work.

### Family Acarnidae

***Megaciella sanctuarium* sp. nov.**

Figures 2, 3, 12

**Figure 12.**
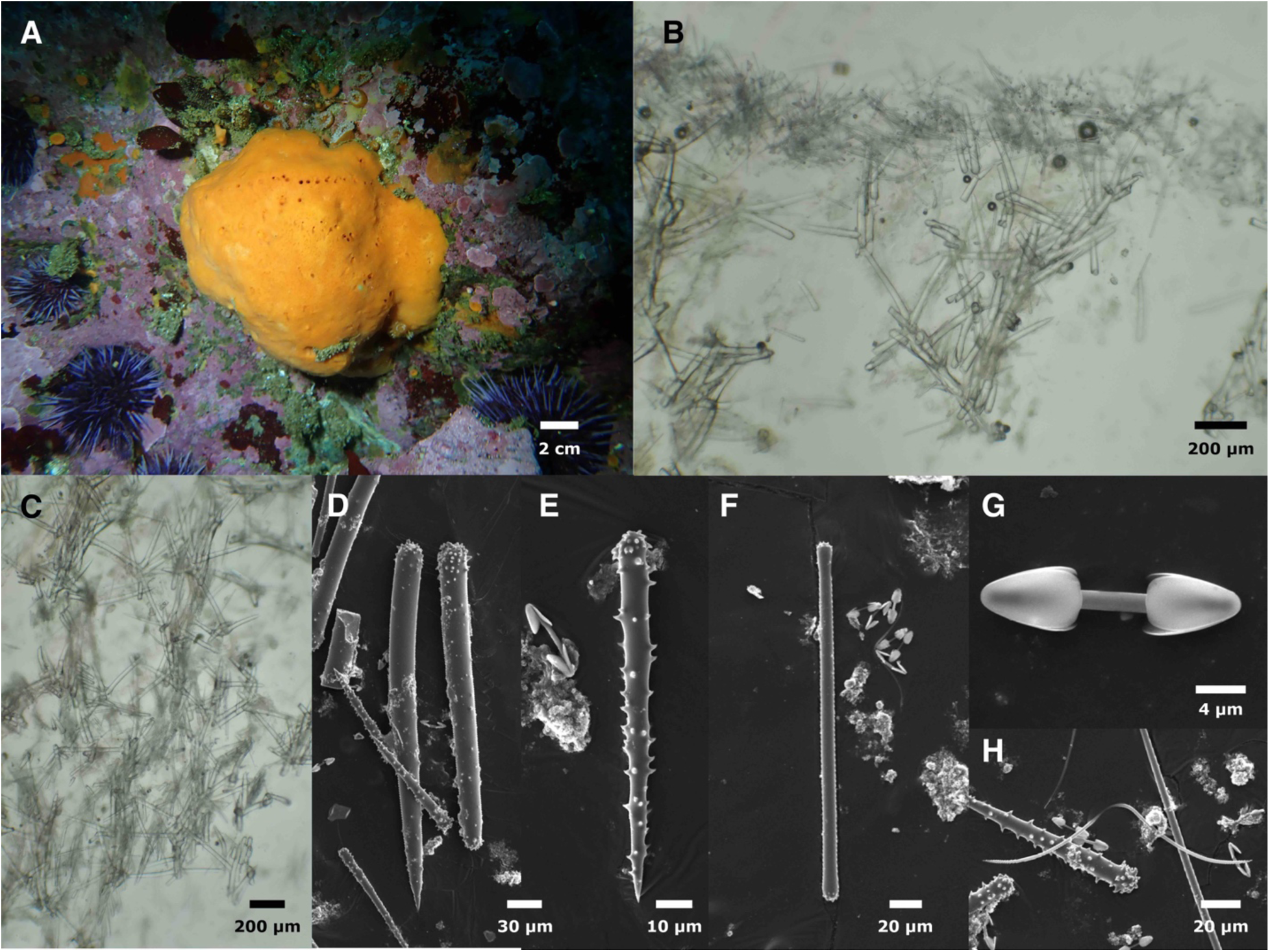
*Megaciella sanctuarium.* A: field photo. B–C: cross sections showing choanosomal skeleton, with ectosomal skeleton visible in B. D: Large acanthostyle and acanthostrongyle. E. small acanthostyle. F: ectosomal strongyle. G: chela. H: Toxa. All images of holotype.

**Material examined.** Holotype: CASIZ236658/IZC00048457, Inner Carmel Pinnacle, (36.55852, −121.96820), 10–24 m, 9/22/21.

**Etymology.** Named in honor of Monterey Bay National Marine Sanctuary.

**Morphology.** Thickly encrusting; sample is 3 cm thick, sponge was approximately 5–8 cm thick when entire. Vivid orange when alive, fades to beige in ethanol. Lumpy and irregular, with oscules occurring in approximate lines across the top of the sponge.

**Skeleton.** Ascending paucispicular tracts of spongin cored with large styles, echinated by small acanthostyles, and accompanied by bundles of thin subtylotes. A chaotic paucispicular reticulation of large styles encased in spongin connects primary tracts. Ectosome a dense mass of thin strongyles/subtylotes, tangential and upright at all angles. Chelae present throughout but most dense at ectosome.

**Spicules.** Large acanthostyles, acanthostrongyles, small acanthostyles, ectosomal strongyles/subtylotes, palmate isochelae, toxas.

Large acanthostyles: small spines concentrated at head, with a few on shaft. 180–302–382 x 8– 20–26 μm (n=25).

Acanthostrongyles: small spines concentrated at each end, with a few on shaft. Some are symmetrical, while others taper slightly towards one end. Appear to be modified large acanthostyles, but slightly shorter. Less common than large acanthostyles. 181–230–289 x 20– 25–31 μm (n=9).

Small acanthostyles: large spines, most dense on head but also covering shaft, head spines curve towards shaft and shaft spines curve towards head. 87–101–116 x 7–9–11 μm (n=22).

Ectosomal strongyles/subtylotes: thin, straight rods with microspined ends; ends variably non– swollen (strongyles) or slightly swollen (subtylotes). 113–204–252 x 3–6–8 μm (n=20).

Palmate isochelae: 17–20–22 μm (n=31).

Toxas: 35–105–168 μm (n=24).

**Distribution and habitat.** Only sample found to date was from the Carmel Pinnacles.

**Remarks.** This species is in the Acarnidae, rather than the Microcionidae, because it possesses ectosomal tylotes rather than styles (Hooper 2002a). The choanosomal skeleton of ascending paucispicular fibers connected by a chaotic unispicular reticulation is consistent with the genus *Megaciella*. Of the 17 named species of *Megaciella*, 10 occur in the Northeast Pacific: 2 in Mexican waters and 10 from the Aleutians and/or the Russian Pacific. Only one of these, *M. microtoxa* (Dickinson, 1945) has acanthostyles. This species, from the Gulf of California, is differentiated from the new species by having much larger coring styles and smaller chelae.

There are also two undescribed species of *Megaciella* from California mentioned by Lee *et al*. (2007). Both species have acanthostyles, but the spicule dimensions reported for these undescribed samples make them unlikely to be additional samples of *Megaciella sanctuarium*.

Genetic data are unable to support or refute the morphological placement of this species. Both loci support a close relationship to the Microcionidae, though with inconsistent placement between the two loci (figures 2 & 3). No other species of *Megacialla* have data available for comparison, and the few other Acarnidae in the phylogenies are inconsistently placed.

This species cannot be identified in the field due to multiple species of sympatric, thickly encrusting orange sponges in the Microcionidae.

### Family Mycalidae

***Mycale (Mycale)* lobos sp. nov.**

Figure 13

**Figure 13.**
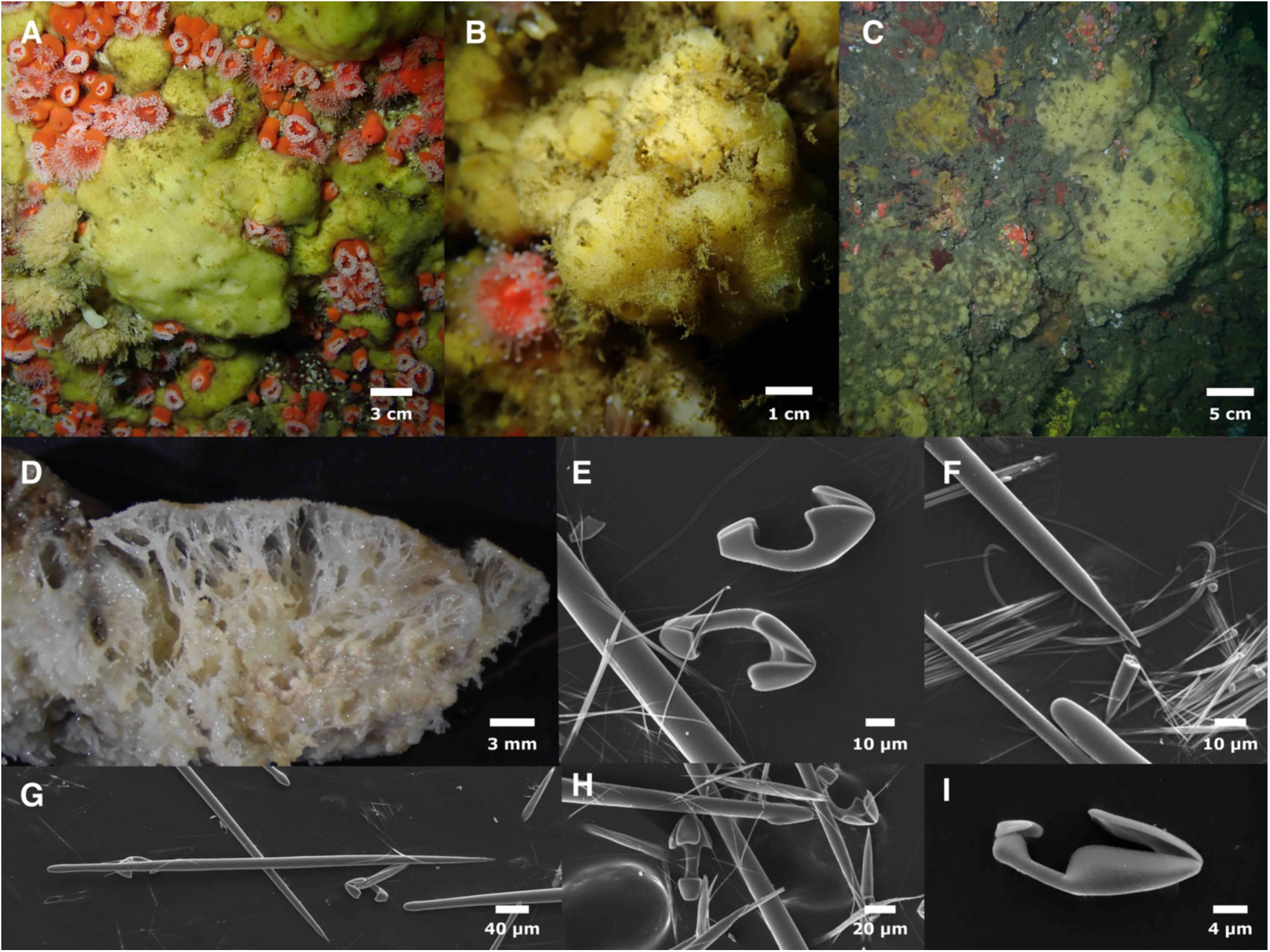
*Mycale lobos.* Field photos of holotype (A), IZC00048458 (B), IZC00048459 (C). D: cross section of SBMNH700920 post-preservation, showing large subectosomal spaces. E, H: Large chelae. F: Sigma. G: style. I: small chela. Raphides visible in E–H. E–I from holotype.

**Material examined.** Holotype: CASIZ236659/IZC00048460, Inner Carmel Pinnacle, (36.55852,-121.96820), 10–24 m, 9/22/21; paratypes: SBMNH700909, Point Pinos, Pacific Grove, (36.64183, −121.93060), 9–18 m, 8/9/21; SBMNH700920, Whaler’s Cove, Point Lobos, Carmel, (36.52172, −121.93894), 6–15 m, 11/23/19; IZC00048458, Dino Head, Point Loma, San Diego, (32.68808, −117.27080), 21–26 m, 9/19/20; IZC00048459, Nudi Wall, Point Loma, San Diego, (32.69872, −117.27580), 18–28 m, 5/16/21, SBMNH700923, Otter Cove, Pacific Grove (36.63110, −121.91840), 9–15 m, 8/9/21.

**Etymology.** Name inspired by Point Lobos, where the first sample was found.

**Morphology.** Thickly encrusting or massive; samples are 2–3 cm thick, but some sponges were up to at least 10 cm thick in life. Irregular in thickness, often appearing as an irregular, featureless blob without visible surface patterning or oscules. Oscules sparse in some samples but more common in others, approximately 2–4 mm in diameter, sometimes found in small clusters. Surface slightly hispid, often with ensnared silt and other debris; lacks surface furrows known to characterize *M. lingua* (Bowerbank, 1866) and *M. toporoki* Koltun, 1958. Ectosome usually pale yellow but sometimes white in life. Large subectosomal vacuoles; in several samples, this space was packed with yellow spheres that appeared to be developing larvae, up to 400 μm in diameter. All tissues white when preserved.

**Skeleton.** Reticulate, anastomosing spongin fibers, cored with styles, rise to the surface of the sponge and support an unstructured mat of styles in the ectosomal skeleton. Large anisochelae occur in rosettes. Trichodramas are extremely abundant.

**Spicules.** Mycalostyles, anisochelae (2 sizes), sigmas (2 sizes), trichodragmas. Mycalostyles: Thickest in center of shaft, but usually subtylote due to constriction (”neck”) below head. Heads generally oblong and sometimes absent; rarely transformed into oxeas. In one sample (IZC00048459) many are polytylote; this was seen rarely in other samples. Significantly shorter in ectosome than choanosome (Wilcoxon rank–sum test p=7.18e–10). Holotype ectosome: 381–471–671 x 7–14–20 μm (n=103). Holotype choanosome: 377–542–631 x 6–13– 18 μm (n=76). If spicule preparations are done from combined ectosomal and choanosomal tissue, length distribution is bimodal, with peaks at ∼450 and ∼550 μm. All samples and tissues pooled: 342–496–671 x 6–13–21 μm (n=401).

Large anisochelae: shaft distinctly curved in profile. Holotype 44–54–63 μm (n=31). All samples pooled 31–51–63 μm (n=158). Shaft length to alae length ratio averages 0.35.

Small anisochelae: Holotype 13–17–24 μm (n=22), all samples pooled 9–17–24 μm (n=169).

Small sigmas: uncommon. All samples pooled 15–18–23 μm (n=5).

Large sigmas: Holotype 45–61–72 μm (n=15), all samples pooled 43–57–72 μm (n=38).

Trichodragmas: abundant. Holotype 33–62–79 x 6–10–13 μm (n=19), all samples pooled 31–58– 76 x 4–9–14 μm (n=28), individual raphides 1–2 μm in width.

**Distribution and habitat.** This species is common but patchily–distributed on subtidal rocky reefs around the Monterey Peninsula (Central California), where it was seen at 5 of 13 reefs investigated. It was also abundant at two sites around the Point Loma kelp forests (San Diego, Southern California), but not seen at the other 6 reefs investigated in that area. Both Point Loma sites were among the deepest sites investigated, with maximum depths of 26–28 m. We therefore hypothesize that it is common in Central California on shallow subtidal reefs, and also common in Southern California, but at slightly deeper reefs.

**Remarks.** The ectosomal skeleton and the lack of modified chelae serve to place this species in the subgenus *Mycale* (*Mycale*). Though there are many species of *Mycale* known from the Eastern Pacific, the curved shaft of the large anisochelae differentiate this species from all but two (Carballo & Cruz-Barraza 2010). *Mycale darwini* Hajdu & Desqueyroux-Faúndez 1994, known from the Galapagos Islands, has curved anisochelae, but grows as a thin crust that is a poor match to the current species. *Mycale toporoki* Koltun, 1958 is more similar. This species was described from Northern Russia, and may also occur in the Aleutian Islands, Pacific Northwest, and possibly California (Lee *et al*. 2007; Lehnert *et al*. 2006). A similar sponge from the North Atlantic, *M. lingua* (Bowerbank, 1866), has also been reported in the North Pacific by some authors, though others have suspected these reports actually refer to *M. toporoki* (Bakus 1966; Lee *et al*. 2007; Lehnert *et al*. 2006). Both *M. lingua* and *M. toporoki* are characterized by surface grooves that are “obvious in living specimens” (van Soest & Hajdu 2002). All sponges of the new species were examined alive and nothing resembling these grooves were seen in any of the specimens. Moreover, the sizes of the styles and both classes of chelae of the new species are much smaller than *M. toporoki* and *M. lingua*. Reports for both of these named species have the smaller size class of chelae as 20–30 μm and the larger size class as 60–100 μm. *Mycale lobos* sp. nov., in contrast, has chelae 11–24 μm and 31–63 μm. Despite measuring 327 chelae, and extensively searching for larger spicules, no chelae longer than 63 μm could be found. No styles longer than 622 μm could be found in the new species either, while the two previously described species have styles up to 800 μm or over 1000 μm, respectively.

This species is not shown in the phylogenies, as we failed to generate genetic data for this species or the congeneric *M. psila* below; this is perhaps due to primer mismatches.

Many specimens of *M. lobos* sp. nov. can be tentatively identified in the field based on gross morphology, albeit mostly by the lack of features. Sponges growing as large, featureless blobs, pale yellow in color, with hispidity collecting debris have thus far proved to be this species upon examination. Tearing the sponge in the field, to look for large cavities filled with developing larvae, would increase confidence of field identification.

***Mycale (Paresperella) psila* (de Laubenfels, 1930)**

Figure 14

**Figure 14.**
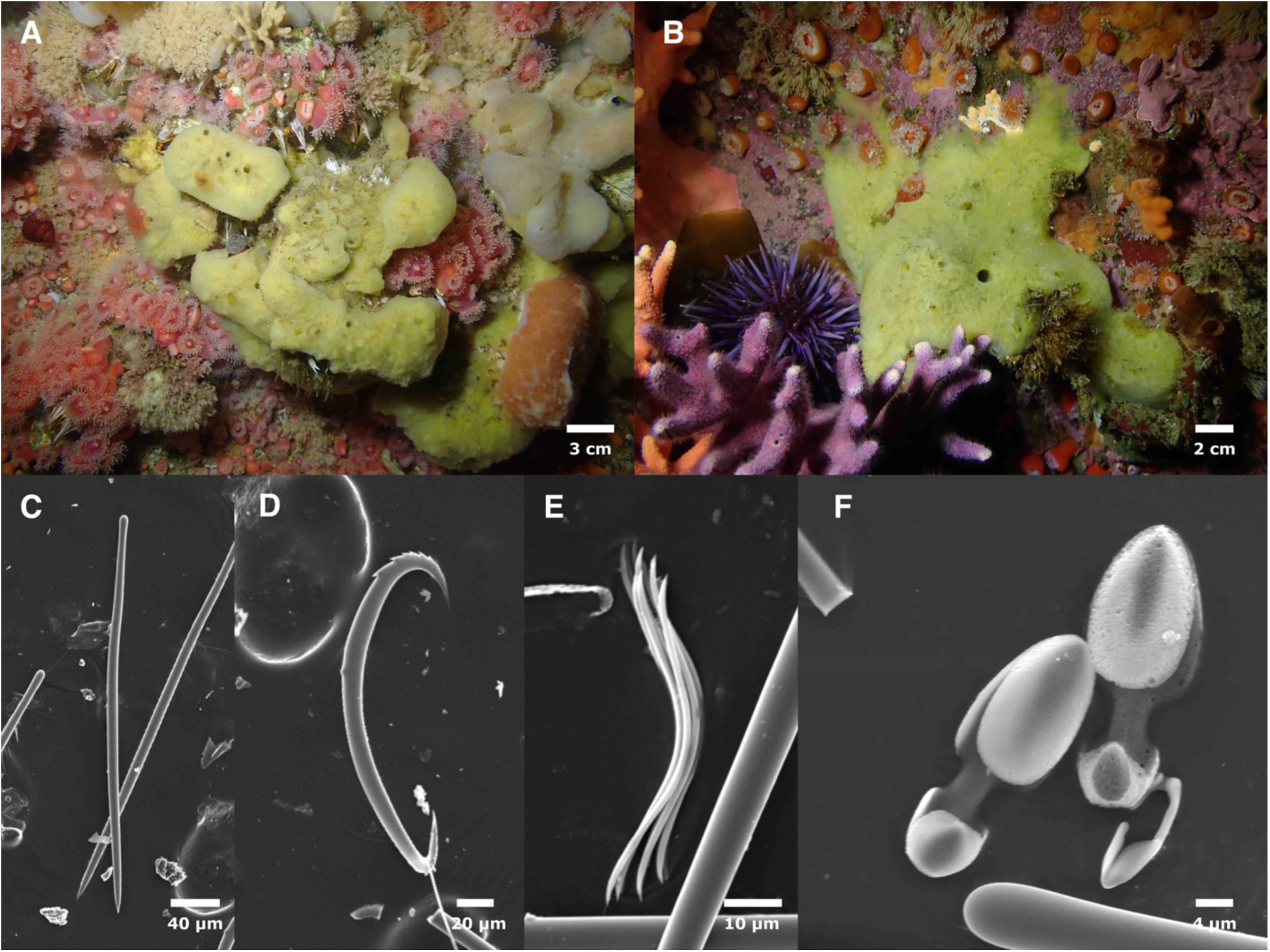
*Mycale psila.* Field photos of IZC00048461 (A) and SBMNH700916 (B). C: style. D: large serrated sigma. E: toxas. F: Large and small chelae. C–F from SBMNH700916.

**Synonymy.**

*Paresperella psila* (de Laubenfels 1930, 1932)

**Material examined.** IZC00048461, Inner Carmel Pinnacle, (36.55910, −121.96630), 10–18 m, 8/10/21; SBMNH700916, Inner Carmel Pinnacle, (36.55852,-121.96820), 10–24 m, 9/22/21.

**Morphology.** Thickly encrusting and forming irregular lobes; samples 1–3 cm thick but sponges were thicker in life; previously reported up to 10 cm thick (Lee *et al*. 2007). Oscula apparent but few in number and chaotically placed; pale yellow in life, white when preserved.

**Skeleton.** Not investigated. Previously reported as having reticulate, anastomosing fibers (Lee *et al*. 2007).

**Spicules.** Mycalostyles, anisochelae (two sizes), sigmas (three types), toxas.

Mycalostyles: slightly sinuous; weakly constricted necks with weak subtylote heads. Sizes not significantly different in ectosome vs. choanosome. 279–336–375 μm (n=75).

Large anisochelae: 27–31–35 μm (n=33).

Small anisochelae: 10–13–19 μm (n=29).

Serrated sigmas: common; C-shaped, thicker than other sigmas and bearing obvious serrations. 110–144–163 μm (n=64).

Large unserrated sigmas: thinner than serrated sigmas, C-shaped in a single plane or S–shaped and twisted so that they are not in a single plane. 126–144–165 μm (n=16).

Small sigmas: C-shaped or shaped like fishing hooks. 34–45–67 μm (n=34).

Toxas: uncommon. 29–55–81 μm (n=16).

**Distribution and habitat.** This species was described by de Laubenfels from Monterey Bay based on two samples (de Laubenfels 1930). It was later reported to occur from British Columbia, Canada, to Southern California, and from the intertidal zone to 26 m (Austin 1985; Lee *et al*. 2007). Our recent collections have found it in the intertidal and shallow subtidal in Central California, but only a single sample was found in the subtidal in Southern California, despite a greater search effort in Southern California.

**Remarks.** The large serrated sigmas serve to differentiate this species from all other sympatric *Mycale*. This species cannot be identified in the field; it resembles other pale-yellow sponges such as *Halichondria loma* sp. nov..

### Order Haplosclerida

**Figure 15.**
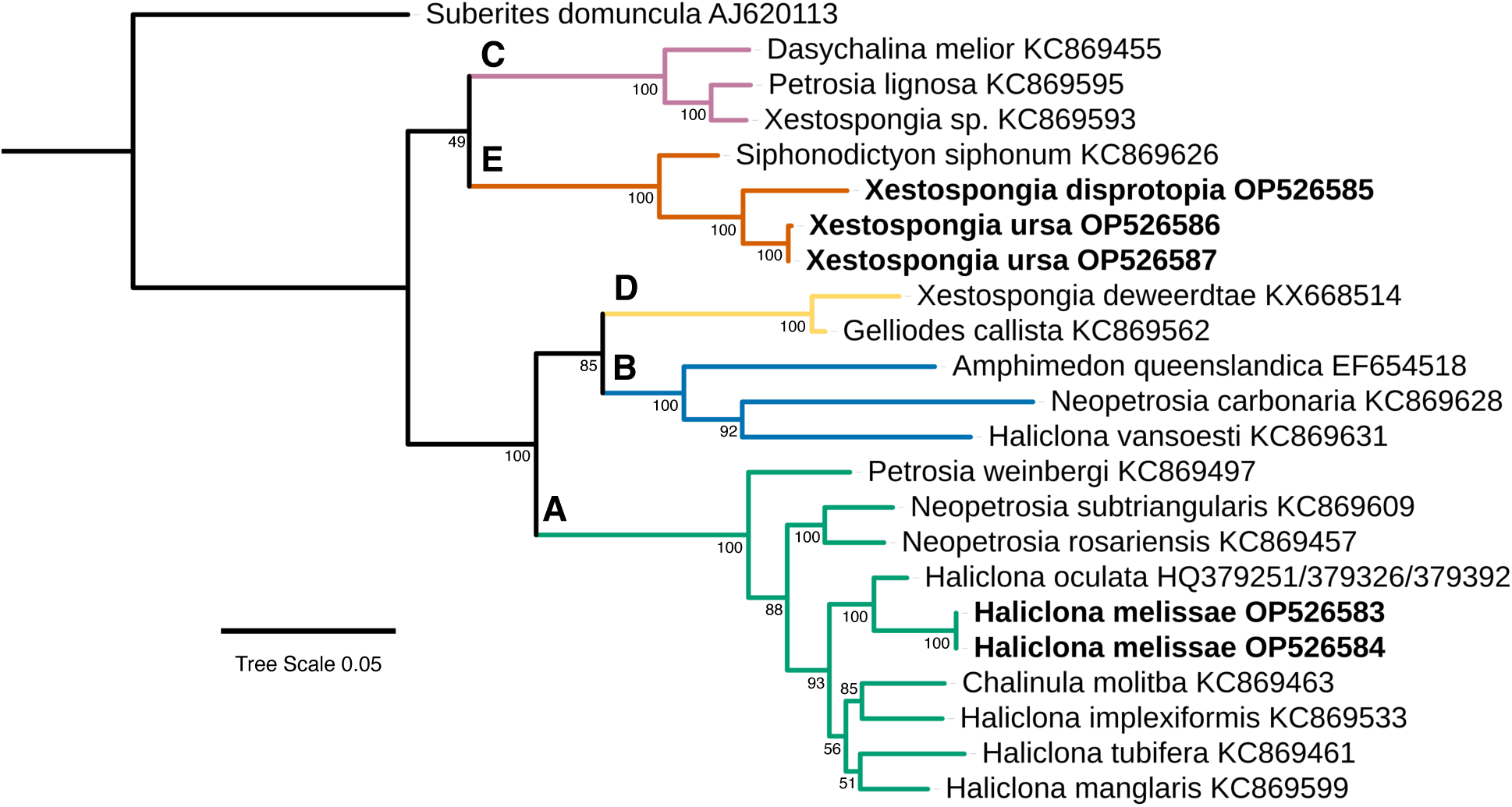
Maximum likelihood phylogeny of the *28S* locus for the Haplosclerida. Colors indicate well-supported clades, with letter names following Redmond *et al*. (2013). Genbank accession numbers are shown; bold indicates new sequences. Node confidence is based on bootstrapping. Scale bar indicates substitutions per site. Colors indicate clades containing new taxa, as referenced in the text.

### Family Petrosiidae

***Xestospongia ursa* sp. nov.**

Figures 15, 16

**Figure 16.**
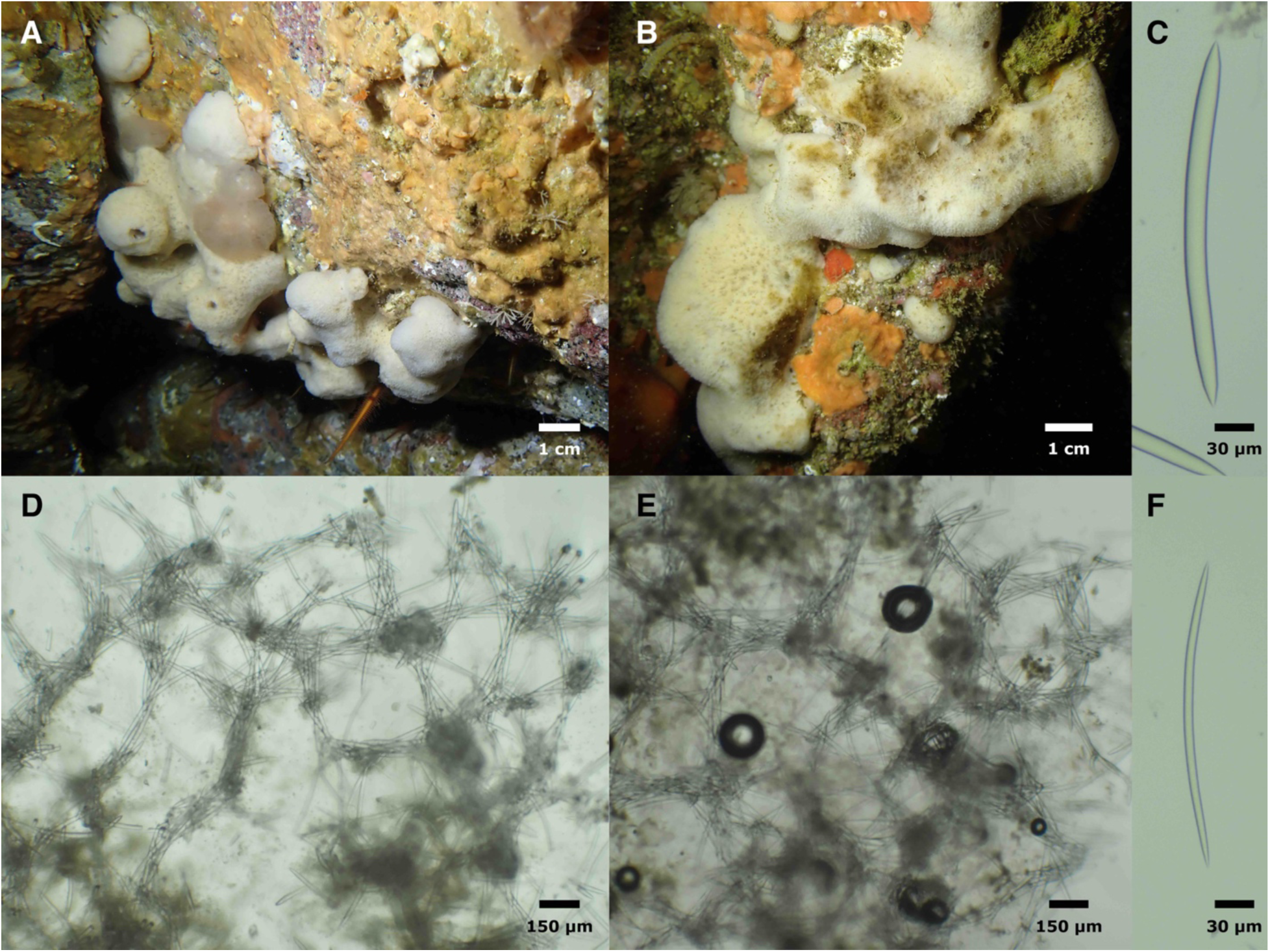
*Xestospongia ursa.* Field photos of holotype (A) and SBMNH700926 (B). C, F: Thick and thin oxeas from IZC00048472. D: Cross-section showing choanosomal skeleton and tangential section showing surface skeleton, both from holotype.

**Material examined.** Holotype: CASIZ236663/IZC00048474, Inner Carmel Pinnacle, (36.55852,-121.96820), 10–24 m, 9/22/21; paratypes: IZC00048472, Fire Rock, Pescadero Point, Carmel, (36.55898, −121.9511), 10–22 m, 8/10/21; SBMNH700926, Tower house arches, Carmel, (36.56187, −121.95950), 9–21 m, 9/21/21; IZC00048473, Whaler’s Cove, Point Lobos, Carmel, (36.52172, −121.93894), 6–15 m, 11/23/19; SBMNH700921, Whaler’s Cove, Point Lobos, Carmel, (36.52172, −121.93894), 6–15 m, 11/23/19. Comparative material: *Xestospongia diprosopia* (de Laubenfels 1932), CASIZ190441 and CASIZ190442, Rittenburg Bank, Farallon Islands, Northern California, 85 m, 10/9/12.

**Etymology.** Bright white and furry-looking, this sponge is named after *Ursa maritimus*.

**Morphology.** Thickly encrusting with irregular lobes and ridges; collected fragments are up to 3 cm thick, but sponges likely exceed this thickness in life. Oscules prominent but sparse and irregular, flush with surface, placed atop high points on lobes and ridges. Surface covered in minute conules that give the sponge a furry appearance. Chalky white alive, but often has brownish patches due to fouling by microscopic algae. Produces copious slime when collected. Very firm and incompressible.

**Skeleton.** Ectosome and choanosome possess a chaotic, reticulated network of single spicules and paucispicular bundles 3–7 spicules across; in some places these form a polygonal mesh. Spongin is not apparent.

**Spicules.** Oxeas, smoothly curved or with one or two subtle bends. Most commonly with two sharp points, but a substantial minority with one end tapering to a blunt tip; a small minority taper at only one end and are therefore styles. Some with extra spines forming a x-shape at one tip; this was very rare in any one sponge, but seen across multiple sponges. Spicules in one length category but variable in thickness; thickness unimodal but some much thinner than average; these may be immature. Holotype 218–290–332 x 4–13–18 μm (n=48), all samples pooled 185–281–332 x 2–14–21 μm (n=201). Mean lengths for each sample vary from 266 to 290 μm.

**Distribution and habitat.** Common on rocky reefs in the shallow subtidal from Pescadero Point, on the northern end of Carmel Bay, to the Carmel Highlands, just south of Carmel Bay; it was seen on six of eight locations in this region. Not seen at any other location to date, including the 6 reefs searched in nearby Monterey Bay.

**Remarks.** *Xestospongia diprosopia* (de Laubenfels, 1930) is a deep-water species from Central California, and the type species for the genus. *Xestospongia ursa* sp. nov. is very similar in terms of skeletal structure, spicule shape, and gross morphology. To verify that *X. ursa* sp. nov. is not a shallow water form of *X. diprosopia*, we examined the spicules of two *X. diprosopia* from 85 m depth. The samples average 351 and 365 μm in length, matching published data from the holotype (352 μm; Desqueyroux-Faúndez & Valentine 2002). This is about 20% longer than *X. ursa* sp. nov., which averages 266–290 μm in length depending on the sponge. The holotype and other *X. diprosopia* also average about 60% thicker spicules (23–26 μm per sample, vs. 13–15 μm for *X. ursa* sp. nov.). We successfully sequenced a portion of the *28S* locus from one *X. diprosopia* sample: this sequence was 6% divergent from *X. ursa* sp. nov. across the aligned 827 bp, supporting the species-level difference of these shallow and deep-water forms (figure 15). The genetic data also support the allocation of the new species to *Xestospongia*, as it is more closely related to the type species than any other species with publicly available data at this locus.

Previous phylogenies of the Haplosclerida have found that most families and genera are polyphyletic, and the position of *X. disprotopia* adds to this conclusion. Most Haplosclerida fall into five clades, designated A–E (Pankey *et al*. 2022; Redmond *et al*. 2013): we include representatives of each in the *28S* phylogeny shown in figure 15. The type species of *Xestospongia* falls within clade E, otherwise populated mainly by members of the Phloeodictyidae. In contrast, other *Xestospongia* have been found to be members of clades B and D (Pankey *et al*. 2022; Redmond *et al*. 2013).

Two other *Xestospongia* are known from the region: *X. dubia* (Ristau, 1978) has much smaller spicules and a different morphology, and *X. hispida* (Ridley & Dendy, 1886) is hispid, with a different morphology and larger spicules.

*Xestospongia ursa* sp. nov. may be identifiable in the field due to its chalky white color, lobe-like growth form, and furry appearance. To date, all samples were correctly identified as conspecifics by the authors before looking at spicules, and no other samples were incorrectly identified as this species among the other samples collected.

### Family Chalinidae

***Haliclona (Halichoclona)* melissae sp. nov.**

Figures 15 & 17

**Figure 17.**
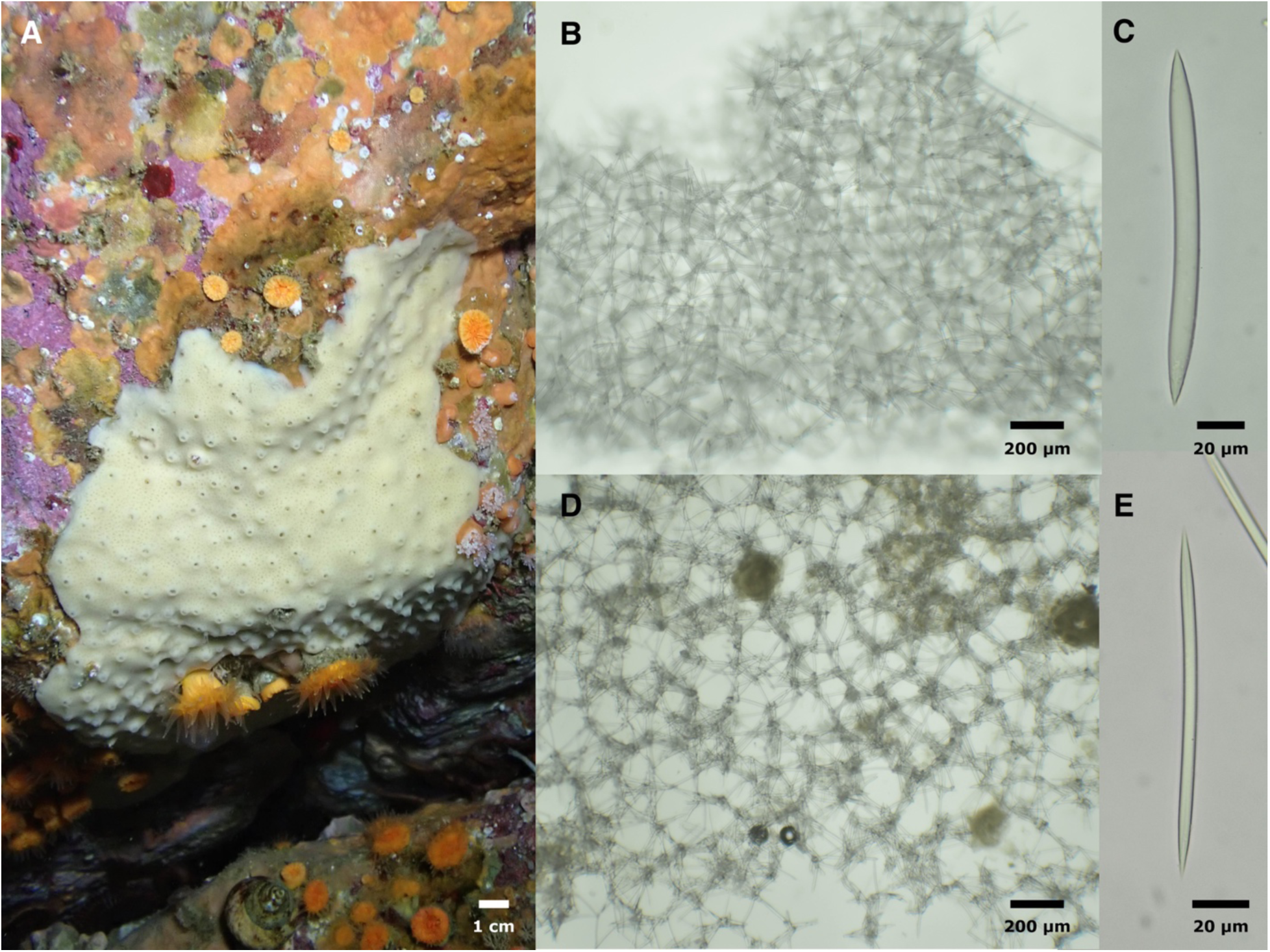
*Haliclona melissae.* A: Field photo. B: Choanosomal skeleton. C: Thick oxea. D: Surface skeleton. E: Thin oxea. All images of holotype.

**Material examined.** Holotype: CASIZ236655/IZC00048448, Inner Carmel Pinnacle, (36.55852,-121.96820), 10–24 m, 9/22/21; paratypes: SBMNH700919 and IZC00048446, Whaler’s Cove, Point Lobos, Carmel, (36.52172, −121.93894), 6–15 m, 11/23/19; SBMNH700927, Wycoff Ledge, San Miguel Island, (34.02132, −120.38710), 9–19 m, 8/25/19; IZC00048447, Tower house arches, Carmel, (36.56187, −121.95950), 9–21 m, 9/21/21.

**Etymology.** Named after Melissa Kamen, a close acquaintance of the first author.

**Morphology.** Encrusting sheets 3–4 mm thick; firm and barely compressible — not spongy as in many *Haliclona* (*Reniera*), nor rock hard as in many Petrosiidae. Surface patterning very regular, oscules approximately 1 mm in diameter, on raised mounds approximately 1 mm in height, each 4–6 mm from next closest oscule; remaining surface patterned with mesh of visible pores. Beige in life and preserved.

**Skeleton.** Ectosome and choanosome comprised of an isotropic, unispicular reticulation with spongin at the nodes.

**Spicules.** Oxeas: more variable in width than length; thin oxeas are possibly immature. A small minority modified to styles. Holotype 108–148–162 x 2–9–12 μm (n=36), all samples pooled: 103–147–168 x 2–9–12 μm (n=146). Mean lengths per sponge 142–154 μm.

**Distribution and habitat.** Found at 3 of 8 shallow subtidal reefs searched from Pescadero Point, on the northern end of Carmel Bay, to the Carmel Highlands, just south of Carmel Bay, but none of the 6 locations searched in Monterey Bay. One additional sample was found at a subtidal site at San Miguel Island in Southern California; San Miguel Island is the outermost of the Channel Islands, with colder waters compared to other Southern California locations.

**Remarks.** The somewhat chaotic nature of the spicule reticulation, the incompressible nature of the sponge, and the hastate oxeas make this species a better match for *Haliclona* (*Halichoclona*) than *Haliclona* (*Reniera*). The skeleton is very similar to *Haliclona* (*Halichoclona*) *gellindra* (de Laubenfels, 1932), the type species for the subgenus, whose type location is the intertidal zone in Carmel Bay. However, the oxeas of the new species are very consistently sized, with mean lengths varying from only 142–154 μm, while spicules in *H. gellindra* are much shorter and thinner: 105–112–122 × 6–6–7 (De Weerdt 2002). The new species is well differentiated from other named *Haliclona* in the region due to skeletal structure, spicule size, and/or the presence of additional types of spicules (Lee *et al*. 2007). There are undescribed species of *Haliclona* that have been noted in previous surveys; where sufficient information is provided to make good comparisons, they match the new species poorly. Previously sequenced species of *Haliclona* have been found in clades A, B, and C, with the largest number in clade A, where the new species is also found.

This species can be tentatively identified in the field in at least some habitats. Samples of this species collected to date are very consistent in form, such that they were correctly identified by the experienced collector before being examined in the lab. A common intertidal *Haliclona* in California, *Haliclona* (*Haliclona*) sp. A (Hartman 1975; Lee *et al*. 2007), has a very similar gross morphology but much smaller spicules and a ladder-like skeleton. *Haliclona melissae* sp. nov. has not yet been found in the intertidal, but if it occurs there, it would likely be confused with this other species without spicule or DNA data. Other undescribed *Haliclona* occur in the subtidal, but those discovered thus far can be differentiated in the field because they are darker in color and much more compressible.

### Order Suberitida

Family Halichondriidae

**Figure 18.**
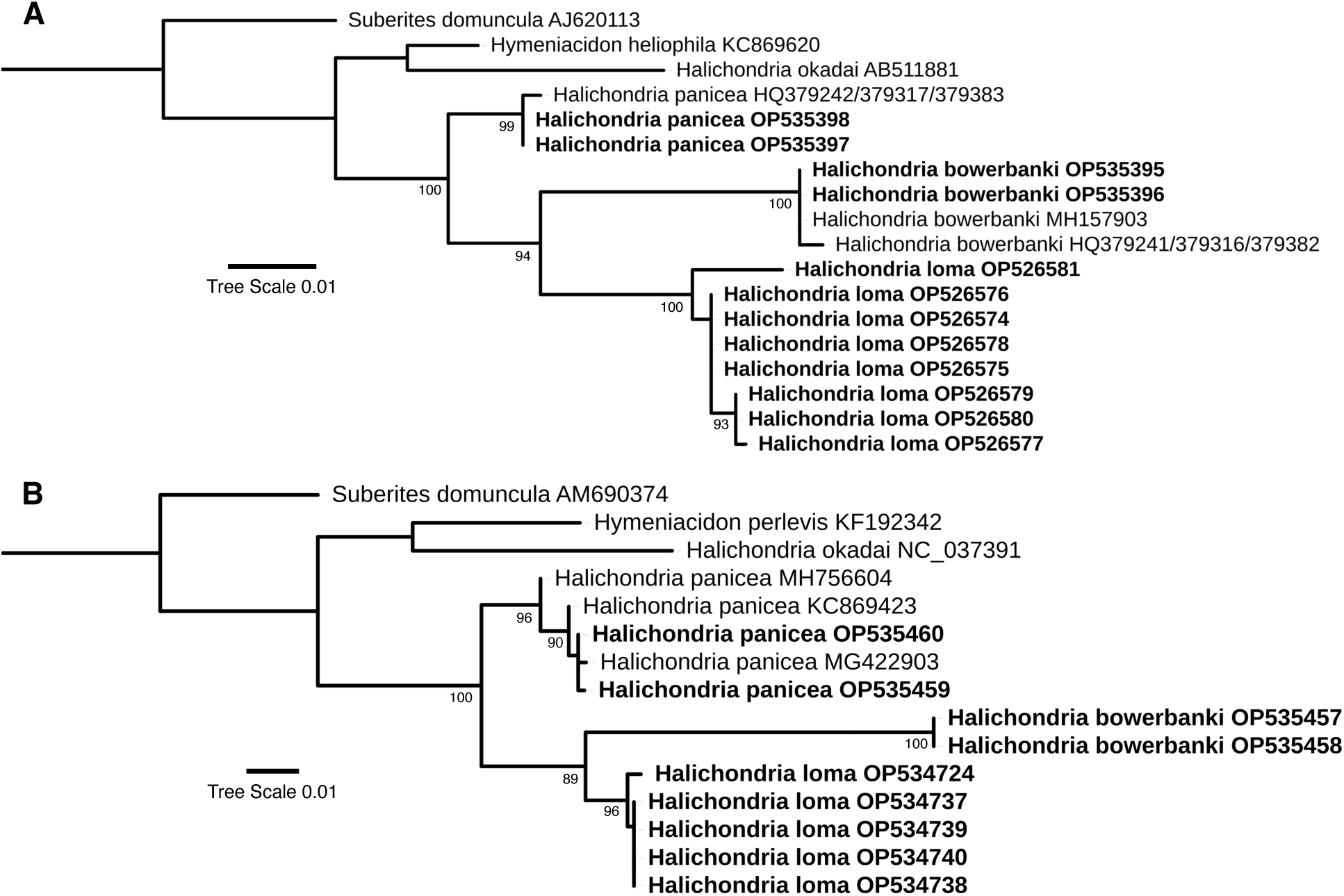
Maximum likelihood phylogenies for the Halichondriidae. A: *28S* locus, B: *cox1* locus. Genbank accession numbers are shown; bold indicates new sequences. Node confidence is based on bootstrapping. Scale bar indicates substitutions per site. Colors indicate clades containing new taxa, as referenced in the text.

***Halichondria loma* sp. nov.**

Figures 18, 19

**Figure 19.**
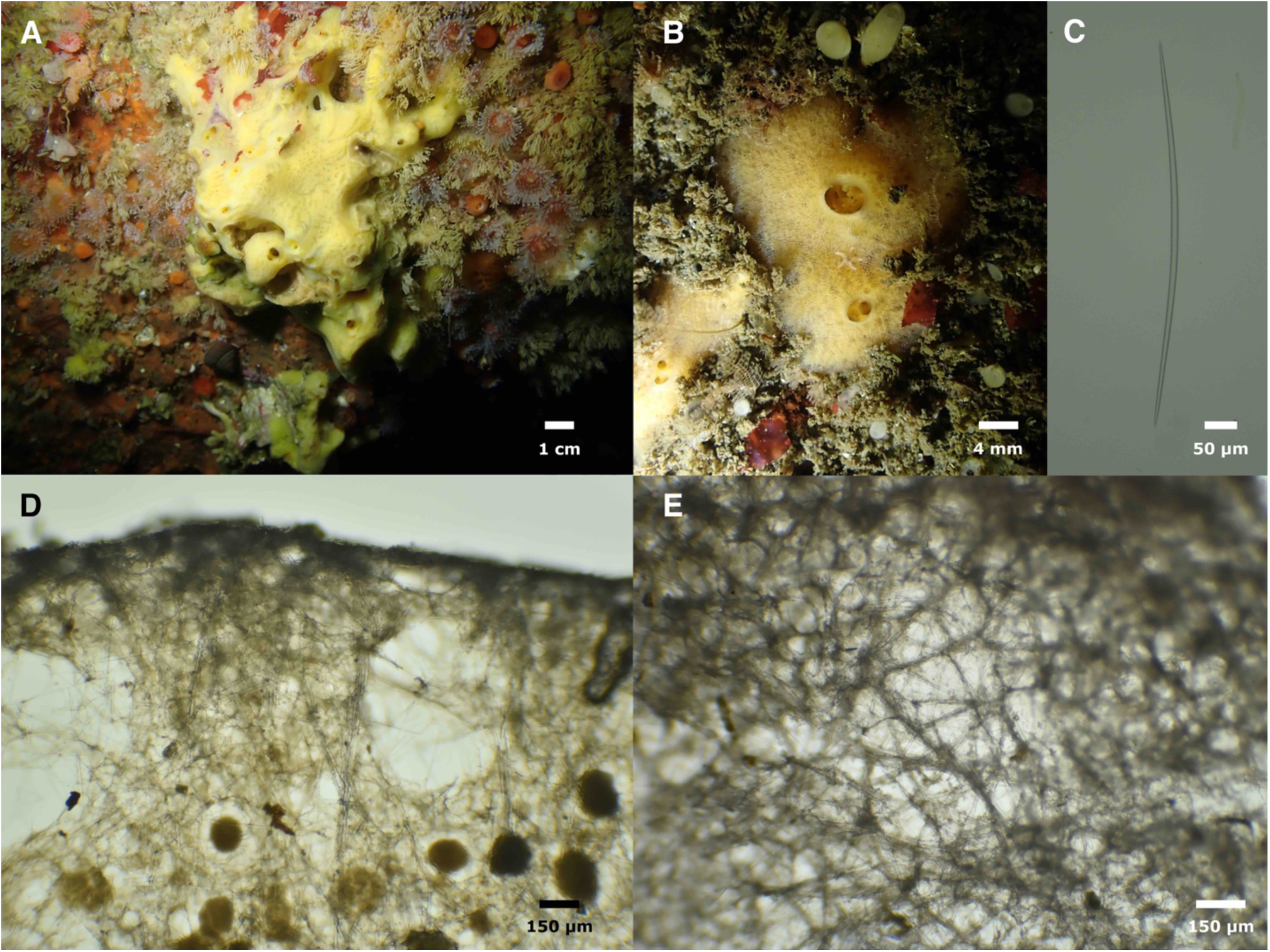
*Halichondria loma.* Field photos of holotype (A) and SBMNH700925 (B). C: oxea from IZC00048442. D: cross-section and E: surface skeleton, both from SBMNH700914.

**Material examined.** Holotype: CASIZ236654/IZC00048445, North Monastery Beach, Carmel, (36.52647, −121.92730), 12–29 m, 9/22/21; paratypes: IZC00048444, Butterfly House, Carmel, (36.53908, −121.93520), 9–20 m, 8/10/21; SBMNH700908, Acropolis Street, Pacific Grove, (36.64183, −121.93060), 9–18 m, 8/9/21; SBMNH700913, Fire Rock, Pescadero Point, Carmel, (36.55898, −121.95110), 10–22 m, 8/10/21; IZC00048442, Inner Carmel Pinnacle, (36.55910, − 121.96630), 10–18 m, 8/10/21; IZC00048443, Wreck of the Ruby E, San Diego, (32.76680, − 117.27620), 18–25 m, 5/16/21; SBMNH700925, Six Fathoms, San Diego, (32.71000, − 117.26860), 9–18 m, 5/15/21; SBMNH700914, Goalpost, San Diego, (32.69438, −117.26860), 12–15 m, 2/8/20.

**Etymology.** Name inspired by Point Loma, the location where the first sample was found.

**Morphology.** Encrusting, bright yellow in life, white when preserved. Oscules prominent, flush with surface or atop low chimneys. Surface patterned with partially transparent mesh-like ectosomal network of fibers and vein-like sub-surface channels. Morphology is very similar to *H. panicea* and *H. bowerbanki*, but in contrast to these species, no individuals were found to form tendrils or occur in green.

**Skeleton.** Choanosome is a chaotic mass of oxeas in confusion, accompanied by paucispicular tracts; no apparent spongin. Ectosomal skeleton of tangential oxeas; in some regions of the sponge, these form a lattice of meandering, paucispicular tracts; in other regions the oxeas form a mat patterned only by circular gaps around pores.

**Spicules.** Oxeas of typical *Halichondria* form, thickest in the center, gradually tapering to hastate tips. Holotype 411–544–650 x 9–12–15 μm (n=40). Other samples average from 462 to 527 μm. All samples pooled: 298–509–650 x 5–11–16 μm (n=386). Only one size class of spicules present (length and width distributions are unimodal), but if spicule preparations are done for ectosome and choanosome separately, spicules are significantly shorter in ectosome (sample IZC00048442: ectosome 409–516–611 x 5–11–15 μm (n=49), choanosome 407–548– 644 x 5–10–14 μm (n=50), Wilcoxon rank-sum test p=0.004).

**Distribution and habitat.** This species is uncommon but widespread on subtidal, natural reefs in Southern and Central California. It was found at 40% of the natural reefs investigated around the Monterey Peninsula, 40% of the natural reefs investigated around San Diego County in extreme Southern California, but not found at any sites between. It was also found on the wreck of the Ruby E, an artificial reef in the San Diego area. It was not common at any site where it was found.

*Halichondria* — *H. panicea, H. bowerbanki*, and/or undescribed species — are abundant in marinas and bays in California, but *H. loma* sp. nov. was not found at any of the 23 marinas investigated by the authors. There are also many intertidal *Halichondria* in Northern and Central California, but *H. loma* sp. nov. was not found at any intertidal sites investigated. *Halichondria loma* sp. nov. therefore seems to be limited to the subtidal, and occurs at low density, primarily on natural reefs.

**Remarks.** Two species of *Halichondria* are known in California: *H. panicea* and *H. bowerbanki*, and the new species is quite similar to both. Previous descriptions of these species in California (de Laubenfels 1932; Lee *et al*. 2007) describe oxeas up to a maximum of 420 μm in length, while the new species is longer: the holotype averages 544 μm, with a maximum of 650 μm. This is a subtle difference, especially considering that *H. bowerbanki* in the United Kingdom has been described as having spicules of similar length (Ackers *et al*. 2007). However, genetic data at mitochondrial and nuclear loci support the distinctiveness of the new species, with *H. panicea, H. bowerbanki*, and the new species all having 14–23% sequence divergence at *28S* and 4–10% sequence divergence at *cox1* (figure 18). The new species is therefore best identified using a combination of genetic and morphological data.

We have extensive collections of *Halichondria* from other locations in Central and Southern California, and these unpublished data support the existence of *H. panicea* and *H. bowerbanki* in the region; in addition to *H. loma* sp. nov., these data point to several other wide-spread undescribed species. These data will be detailed in an upcoming systematic revision of the family Halichondriidae for the region.

This species cannot be identified in the field, as the gross morphology cannot be differentiated from *H. panicea* or *H. bowerbanki. Halichondria* can be tentatively identified to the genus level in the field, but are easy to confuse with some other species, such as *Mycale psila* and *Hymeniacidon perlevis*.

***Hymeniacidon fusiformis* sp. nov.**

Figures 18 & 20

**Figure 20.**
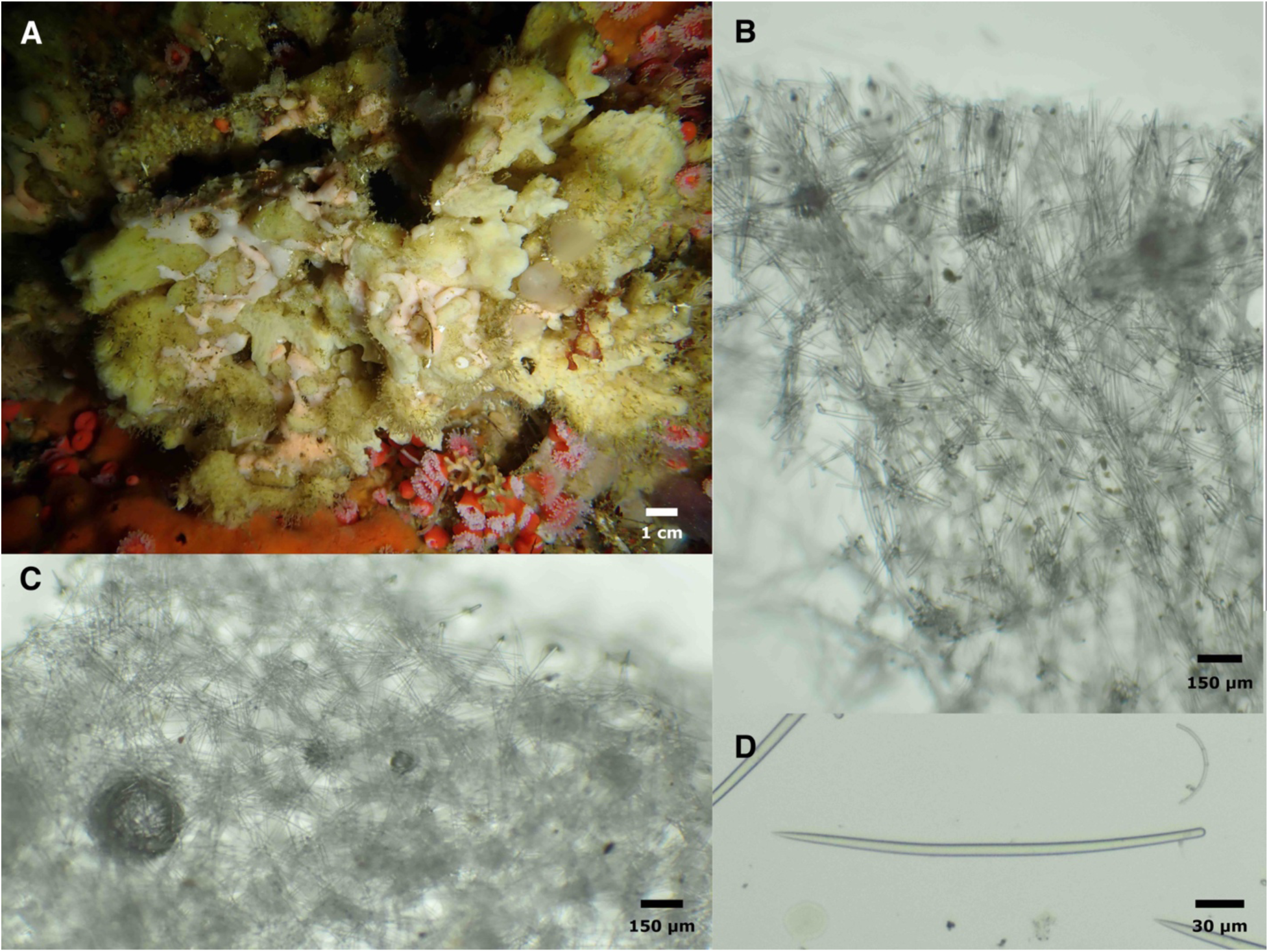
*Hymeniacidon fusiformis.* A: field photo. B: cross-section at sponge surface. C: ectosomal skeleton. D: oxea. All images of holotype.

**Material examined.** Holotype: CASIZ236657/IZC00048452, Inner Carmel Pinnacle, (36.55852, −121.96820), 10–24 m, 9/22/21.

**Etymology.** Named for its slightly fusiform styles.

**Morphology.** Growing as an irregular mass, thickly encrusting with many projecting fingers and lobes, heavily fouled by hydroids and other animals. Sampled portion is 1 cm thick, but sponge was thicker in life. Firm and barely compressible; oscules and pores not evident.

**Skeleton.** Choanosome a chaotic halichondrid reticulation with vague multispicular tracts meandering towards the surface. Spicule density high; spongin not apparent. Ectosome a dense mat of tangential styles approximately 200 μm thick, unstructured in some regions, and patterned into a semi-regular mesh in other regions. Large subectosomal spaces present, bridged by spicule tracts to support ectosome.

**Spicules.** Styles: most are very slightly fusiform, so that they are thicker in the center than near the head of style. Length and width are both highly variable; distributions are continuous but bimodal, 186–281–409 x 4–10–18 μm (n=345), length modes 260 and 330 μm, width modes 8 and 13 μm. When spicule preps are done separately for ectosome and choanosome, no differences are found in spicule sizes by domain: ectosome 217–286–372 x 4–11–18 μm (n=68), choanosome 206–285–370 x 6–10–18 μm (n=62), Wilcoxon rank-sum test p = 0.863.

**Distribution and habitat.** Known only from the Carmel Pinnacles.

**Remarks.** Of the three *Hymeniacidon* known from the region, *H. actites* (Ristau, 1978) and *H. ungodon* de Laubenfels 1932 have much smaller spicules, with less variation in spicule dimensions. *Hymeniacidon perlevis* (Montagu, 1818) have spicules of similar length, and also have high variation in spicule length. However, *H. perlevis* spicules vary little in width and are not fusiform. The new species also differs from *H. perlevis* in coloration: though *H. perlevis* can be yellow, orange, or red, it is not known to occur in white. Finally, *H. perlevis* spicules differ in having a unimodal distribution for both length and width, and being significantly smaller in the ectosome than the choanosome. These species are also genetically differentiated at the *28S* locus, as shown in figure 18. It is unlikely that this species can be identified in the field.

### Order Scopalinida

#### Family Scopalinidae

**Figure 21.**
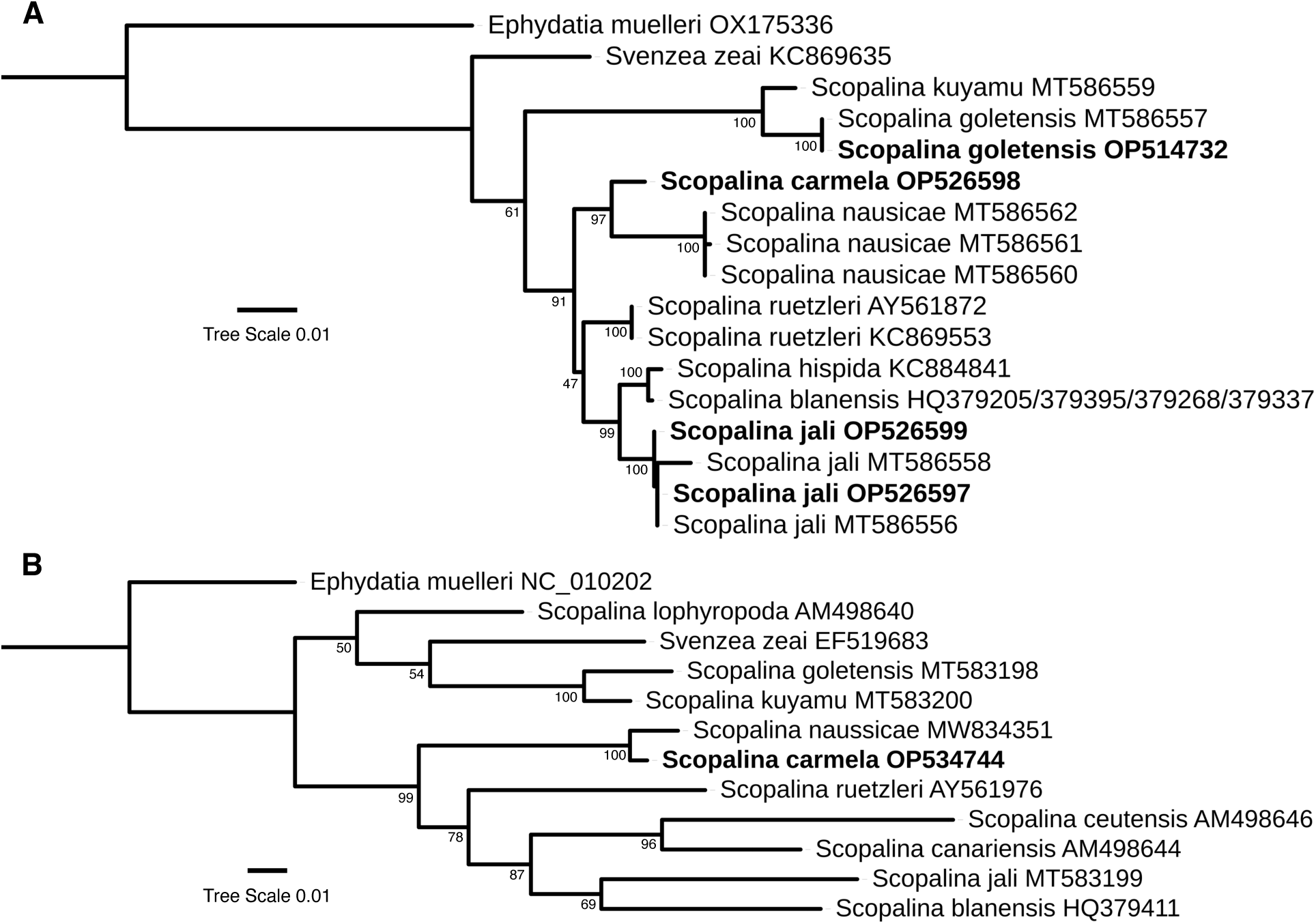
Maximum likelihood phylogenies for the Scopalinida. A: *28S* locus, B: *cox1* locus. Genbank accession numbers are shown; bold indicates new sequences. Node confidence is based on bootstrapping. Scale bar indicates substitutions per site. Colors indicate clades containing new taxa, as referenced in the text.

***Scopalina carmela* sp. nov.**

Figures 21 & 22

**Figure 22.**
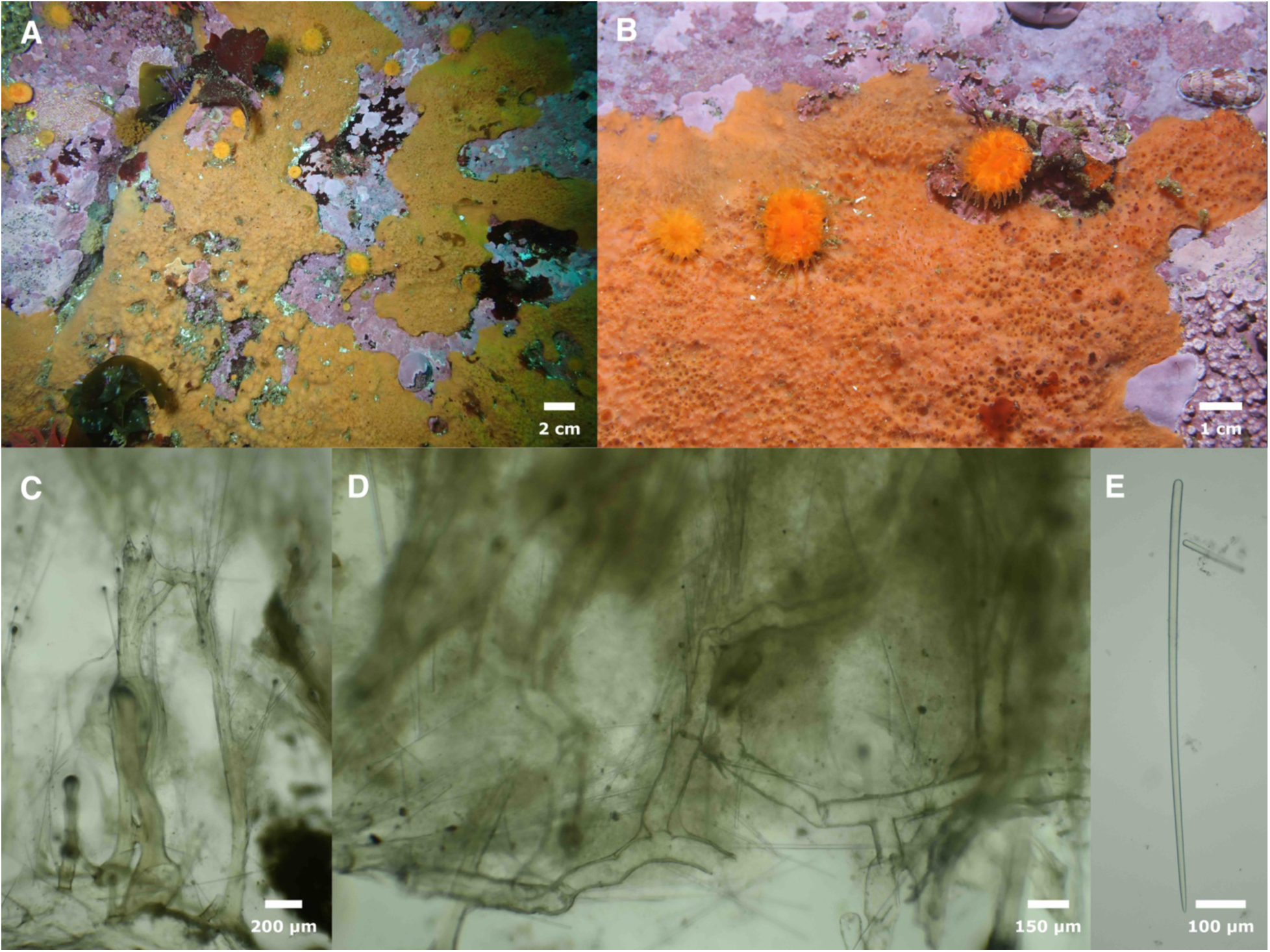
*Scopalina carmela.* Field photos of holotype (A) and unsampled individual (B). C: cross-section showing spongin plate and vertical nodes, with associated spicules from holotype. D: Cross-section showing vermiform tracts of spongin, from holotype. E: style from SBMNH700917.

**Material examined.** CASIZ236662/IZC00048468 (holotype) and SBMNH700917 (paratype), Inner Carmel Pinnacle (36.55910, −121.96630), 10–18 m, 8/10/2021.

**Etymology.** Named for Carmel Bay.

**Morphology.** Encrusting, 4–6 mm thick. Light orange alive, beige when preserved. Surface is covered in abundant conules, oscula, and large pores.

**Skeleton.** Nodes of spongin arise from a basal spongin plate and terminate in surface conules. Vertical spongin nodes are occasionally bridged by secondary horizontal branches of spongin. Primary and secondary spongin tracts are cored by styles, either entirely enclosed by spongin or with heads embedded and points emerging; styles are chaotically arranged but generally at an angle between vertical and 45 degrees, tips up. Styles are more abundant near the sponge surface, where they form bouquets at the top of primary spongin tracts, piercing the surface at conules. Sponge also contains meandering, vermiform tracts of spongin that are not cored with styles. Sponge contains abundant sand, but no debris was seen coring spongin tracts. Tracts are often filled and/or coated with what appear to be red algal cells.

**Spicules.** Long smooth styles that taper gradually to a sharp point; sometimes with subtle step-changes in size near the tip (”telescoping tips”). Holotype 713–915–1132 x 8–13–16 μm (n=26), other sample: 606–809–1017 x 10–16–21 μm (n=20).

**Distribution and habitat.** Known only from the Carmel Pinnacles.

**Remarks.** This species is clearly within the *Scopalina* based on both morphology and genotype. Four species of *Scopalina* are known from this region, all recently described from Southern California (Turner 2021). The skeleton and general appearance of this new species is very similar to *S. nausicae*, but it is easily differentiated from that species by having spicules nearly twice as long (*S. carmela* sp. nov. mean length = 869, *S. nausicae* mean length = 494), and also by its orange color (*S. nausicae* is peach-colored). Several pieces of evidence support a hypothesis that *S. carmela* sp. nov. is a distinct species, and not a northern form of *S. nausicae.* First, the morphological differences (color and differences in spicule length) are very similar to the differences between *S. kuyamu* and *S. goletensis*, which both occur in Southern California. Second, genetic differentiation between *S. carmela* sp. nov. and *S. nausicae* are of a similar magnitude as the differentiation between *S. kuyamu* and *S. goletensis* at both *cox1* and *28S*. Third, the final California species of *Scopalina, S. jali,* is found at both the Carmel Pinnacles and in Southern California, without genetic differentiation at *28S* (figure 21).

Close examination or high-resolution photos make this species identifiable in the field: the highly conulose surface, abundant pores and oscules, and light orange color serve to distinguish this species from other California species. Other encrusting orange sponges in California are firm, while this species is very soft and compressible.

***Scopalina jali* Turner, 2021**

Figures 1B & 21

**Material examined.** IZC00048470 and SBMNH700918, Inner Carmel Pinnacle (36.55910, − 121.96630), 10–18 m, 8/10/2021.

**Morphology.** Thickly encrusting, 1–2 cm thick, up to 20 cm across. Soft and compressible. Prominent oscules slightly elevated above surface, Ectosome partially transparent and lacy. Terracotta (reddish-brown) to beige alive, beige in ethanol.

**Skeleton.** Not investigated. Previously reported as a chaotic mesh of spongin fibers cored with spicules.

**Spicules.** Oxeas, rarely modified to anisoxeas or with an extra cross-piece forming an X shape at one end. 361–423–485 x 7–15–18 μm (n=22).

**Distribution and habitat.** Occurs on natural reefs in the shallow subtidal. Previously known only from Southern California, from Santa Barbara in the north to Catalina Island in the south. This species was common at both dives at the Carmel Pinnacles, extending its range into Central California. It has not been seen at any of the 13 other shallow subtidal sites recently investigated around the Monterey Peninsula.

**Remarks.** The spicules of the sample investigated are 15% longer and 40% thicker than the Southern California samples measured in previous work (Turner 2021). As no genetic differentiation was seen at the *28S* locus, this is likely due to differences in environmental conditions.

Order Axinellida

Family Raspailiidae

***Endectyon (Endectyon) hyle* (de Laubenfels, 1930)**

Figure 1F.

**Synonymy.**

*Hemectyon hyle* (Bakus & Green 1987; Dickinson 1945; Green & Bakus 1994; de Laubenfels 1932)

*Aulospongus hyle* (Desqueyroux-Faúndez & van Soest 1997)

*Raspailia (Raspaxilla) hyle* (Aguilar-Camacho & Carballo 2013; Hooper *et al*. 1999)

**Material examined.** IZC00041043, Inner Carmel Pinnacle, (36.55852,-121.96820), 10–24 m, 9/22/21.

**Morphology.** Upright, bushy lobes rise from a broad base, expand and occasionally branch; oscules at high points; highly hispid; reddish orange in life, fades to tan in ethanol. Previously reported as occurring in both reddish-orange and yellow (Turner & Pankey 2023).

**Skeleton.** Not investigated. Previously described as a dense, axially-compressed reticulation of styles, echinated by acanthostyles; long subectosmal styles pierce the surface, surrounded by bouquets of thin ectosomal styles (Turner & Pankey 2023).

**Spicules.** Short styles, long styles, acanthostyles, and thin ectosomal styles, rare oxeas.

Short styles: unadorned, often curved or bent, with the bend most commonly near the head end but often in the middle. 393–504–605 x 10–19–25 μm (n=28).

Long styles: unadorned, straight or curved. 1248–1484–1791 x 12–21–27 μm (n=6).

Acanthostyles: slightly curved or straight; large curved spines limited to the half of style closer to the point, curved towards head end of style. 206–316–409 x 15–21–29 μm (n=22).

Thin ectosomal styles: 382–491–611 x 2–3–5 μm (n=14).

Oxeas: one seen, 868 x 24 μm.

**Distribution and habitat.** Known from British Columbia to the Gulf of California, and from 15 m to 330 m in depth (Turner & Pankey 2023)

**Remarks.** This species is included in a recent revision of the order Axinellida in California: see Turner & Pankey (2022) for more details.

### Order Bubarida

**Figure 23.**
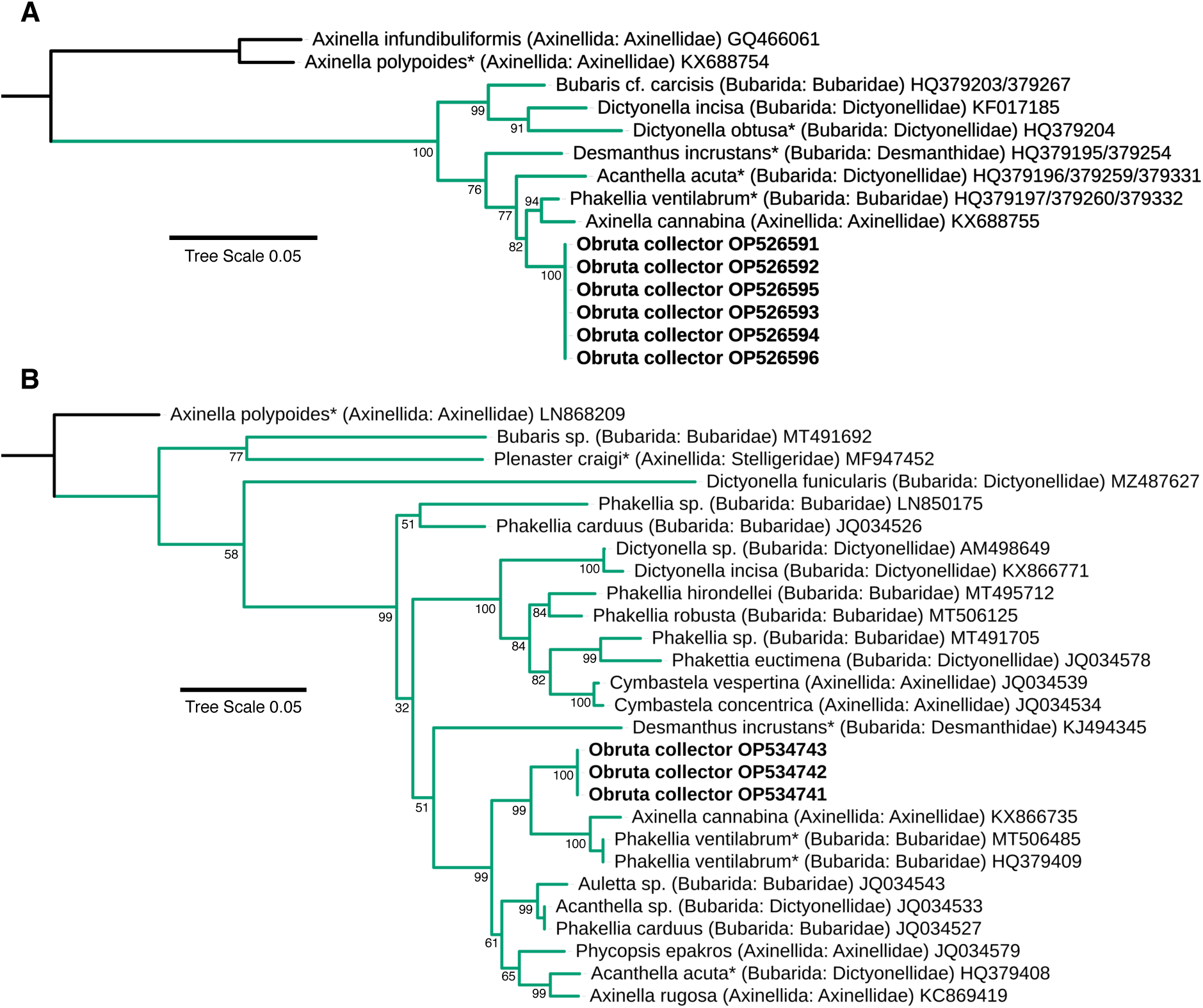
Maximum likelihood phylogenies for the Bubarida. A: *28S* locus, B: *cox1* locus. Green clade designated the putative Bubarida; the orders and families currently housing each taxon is also shown. Genbank accession numbers are shown; bold indicates new sequences; asterisks designate type species. Node confidence is based on bootstrapping. Scale bar indicates substitutions per site. Colors indicate clades containing new taxa, as referenced in the text.

**Remarks.** The order Bubarida is ill-defined and in need of systematic revision. The order was created in response to genetic support for a clade containing species of *Acanthella, Dictyonella, Bubaris, Phakellia* and more (Morrow & Cárdenas 2015). This clade was supported by a recent multi-locus phylogeny (Pankey *et al*. 2022, figure S5); our single-locus phylogenies, which have less resolution but more taxa, are shown in figure 23. Though some species currently placed in the Axinellida are within this green clade, these species are expected to be moved to the Bubarida upon revision, as the type species *Axinella polypoides* is in a separate clade (Gazave *et al*. 2010; Pankey *et al*. 2022; Turner & Pankey 2023).

### Family Bubaridae

*Obruta* gen. nov.

**Type species.** *Obruta collector* sp. nov.

**Diagnosis.** Bubaridae with oxeas as primary spicules, thickly encrusting growth habit, and very hispid surface. Slightly sinuous oxeas found near the sponge base, mixed in confusion with non-sinuous oxeas; ill-defined tracts of non-sinuous oxeas rise vertically from this base.

**Etymology.** Named for the copious debris often trapped by the long protruding spicules. From the latin obruta, meaning debris.

***Obruta collector* sp. nov.**

Figures 23 & 24

**Figure 24.**
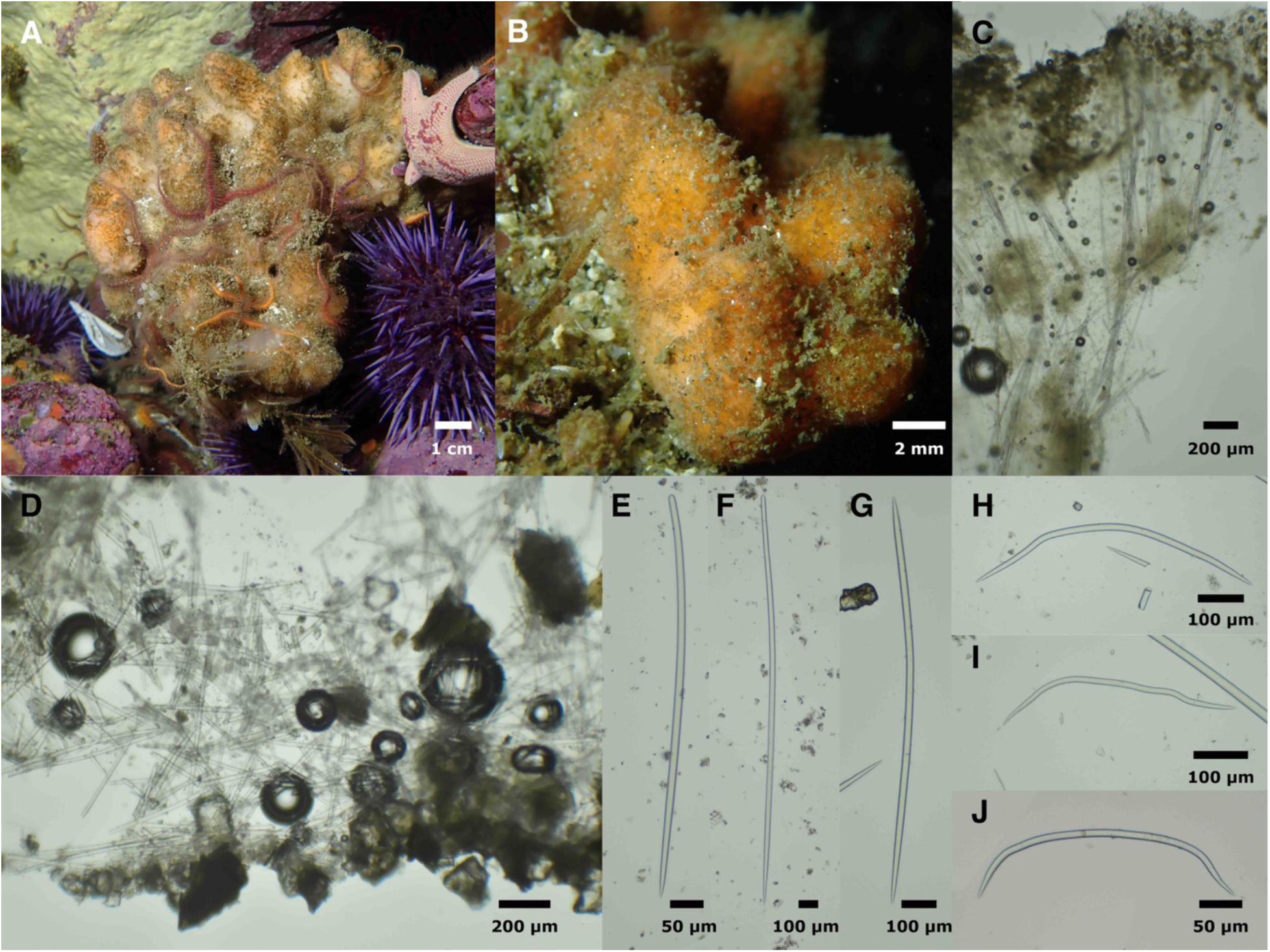
*Obruta collector.* Field photos of IZC00048463 (A) and holotype (B). *Leucilla nuttingi* sample IZC00048456 visible as small calcareous growths on *O. collector* in A. C: cross-section at surface of sponge, from holotype. D: Cross-section at base of sponge, from TLT522.

E: style from SBMNH700907. F: Anisoxea from SBMNH700907. G: oxea from IZC00048464. H–J: sinuous oxeas. H,I from holotype, J from SBMNH700924

**Material examined.** Holotype: CASIZ236660/IZC00048466, La Jolla Cove, San Diego (32.85227, −117.27239), 10–16 m, 8/14/20. Paratypes: Goalpost Reef, San Diego (32.69438, − 117.26860), 12–15 m, 2/8/20; IZC00048464, Isla Vista Reef, Santa Barbara (34.40278 − 119.85755), 9–12 m, 8/1/19; IZC00048462, Arroyo Quemado Reef, Santa Barbara (34.46775, − 120.11905), 7–11 m, 6/14/19; SBMNH700907, Arroyo Quemado Reef, Santa Barbara (34.46775, −120.11905), 7–11 m, 7/29/19; IZC00048465, Whaler’s Cove, Point Lobos, Carmel (36.52172, −121.93894), 6–15 m, 11/23/19; SBMNH700924, Otter Cove, Pacific Grove (36.62920, −121.92031), 7–12 m, 11/24/19; IZC00048463, Inner Carmel Pinnacle (36.55852, − 121.96820), 10–24 m, 9/22/21; SBMNH700915, Goalpost, San Diego, (32.69438, −117.26860), 12–15 m, 2/8/20.

**Etymology.** “Debris collector,” because the long protruding spicules tend to collect copious detritus in the habitats where it is found.

**Morphology.** A thickly encrusting sponge with upright lobes. Samples 3 mm to 2 cm thick; incompletely sampled individuals likely exceeded 4 cm in height. Color in life apricot (light yellow-orange); some samples variegated with peach and white; all samples beige when preserved. Surface is thick with protruding spicules, giving the sponge a furry appearance, often with sediment and debris ensnared on hispid surface.

**Skeleton.** Upright, slightly plumose bundles of oxeas, 5–10 spicules wide, project towards sponge surface, where they pierce the tissue and create a hispid surface. Many spicules also found outside of bundles; near sponge surface, these are mostly upright, adding to a generally plumose impression; near the sponge base, oxeas are found in utter confusion. Some megascleres found horizontally where sponge attaches to substrate; a minority of these have multiple bends resulting in a U or S shape and could be dubbed sinuous. No ectosomal specialization. No apparent spongin; spicules are unbound after proteinase K digestion. Spicule density is high; sponges are firm and incompressible.

**Spicules.** Oxeas, styles, and intermediates; sinuous oxeas.

Oxeas: slightly curved or straight; majority with sharp points at both ends, but minority have asymmetrical points, with one tip blunt or rounded; some of these approach styles. Holotype: 502–750-1208 x 5–14–31 μm (n=80). Dimensions vary greatly within and among samples. Average length was significantly associated with latitude (r^2^ = 0.6, p = 0.02), but width was not (r^2^ = 0.4, p = 0.08). Average lengths: San Diego samples = 750 and 747 μm; Santa Barbara samples = 743, 796, and 878 μm; Monterey samples = 932, 1138, and 1439 μm. All samples pooled: 477–948–2155 x 3–15–31 μm (n=360).

Styles: shaped like oxeas, but with one untapered, rounded end; less common than oxeas in all samples, with frequencies of approximately 1%–15%, depending on sample. Shorter than oxeas in all samples, with average lengths 75%–85% as long. Holotype: 521–612–729 x 8–12–15 μm (n=5); all samples pooled: 486–717–1321 x 5–12–23 μm (n=38).

Sinuous oxeas: oxeas with multiple bends and curves such that they are sinuous or U-shaped; some are only slightly sinuous and hard to differentiate from normal oxeas. Found in 5 of 8 samples. Holotype: 371–504–630 x 7–11–16 μm (n=12); all samples pooled: 231–510–1692 x 7–11–18 μm (n=43).

**Distribution and habitat.** Found on shallow subtidal reefs in Central and Southern California; not found in the intertidal or on human structures. Fairly common around Monterey and Carmel Bays, where it was found at 50% of subtidal reefs investigated. Less common in Southern California, where it was found at only 7% of subtidal reefs.

**Remarks.** *Obruta collector* sp. nov. is confidently placed in this nominal Bubarida clade at both loci, forming a subclade with *Phakellia ventilabrum*, type of its genus, and *Axinellia cannabina* (considered by some authors to be an *Acanthella* (Gazave *et al*. 2010)). The close evolutionary relationship of these three species highlights how difficult it will be to revise this order based on traditional characters. The two previously described species possess an axial skeleton of sinuous strongyles complimented by styles, and the new species possesses mostly oxeas. *Phakellia ventilabrum* is upright and fan-shaped, *Axinella cannabina* is branching and tubular, and *Obruta collector* sp. nov. is thickly encrusting and lobed.

The high spicule density, hispid surface, and basal skeleton of sinuous diactines of *O. collector* sp. nov. are consistent with placement in the Bubaridae, and not consistent with the other bubarid families (Dictyonellidae and Desmanthidae). However, none of the existing genera placed within the Bubaridae, nor other genera that may be included in the Bubarida after revision, are a good fit for *O. collector* sp. nov. The close relationship of this new species to the type species of *Phakellia* is intriguing, but that genus is currently defined by its planer habit, multiple axes of sinuous strongyles, and styles, none of which are possessed by *O. collector* sp. nov. The genus *Bubaris* is more similar to the new species, as it is encrusting with sinuous diactines along the substrate. However, *Bubaris* is defined by having sinuous strongyles and styles, a thinly encrusting habit, and a skeleton of single styles erect on the substrate. Though some species currently placed in this genus have other growth forms (e.g., *B. sarayi* (Ilan *et al*. 1994, *B. conulosa* Vacelet & Vasseur 1971, they all possess only sinuous strongyles and styles. As *O. collector* sp. nov. possesses primarily oxeas and is thickly encrusting, we deem it necessary to create a new genus to house this species. The presence of sinuous diactines and lack of an ectosomal skeleton are morphological characters that unite this species with most other members of the Bubarida and other species that will likely be revised into the genus based on the phylogenies shown.

*Obruta collector* sp. nov. can be easily differentiated from other named species, as evidenced by the need to create a new genus to house it. Due to the ill-defined boundaries of some genera of Bubarida, Axinella, and Halichondriidae, however, it is worth listing the similarities and differences of species from several other genera that occur in the North East Pacific. The only previous Bubarida known from the region is *Rhaphoxya laubenfelsi* Dickinson, 1945 from Pacific Mexico. This species is differentiated by being ramose and branching, possessing sharply bent styles, and being green after preservation. Two *Axinyssa* are known from the northeast Pacific, and some members of this genus have genetic affinities to the Bubarida. *Axinyssa tuscara* (Ristau, 1978) is sympatric with *O. collector* sp. nov., but is dark brown with smaller oxeas. Another species from Pacific Mexico, *Axinyssa isabela* Carballo & Cruz-Barraza, 2008, has similar spicules and a somewhat similar skeleton to *O. collector* sp. nov., but is yellow, sprawling, and is not macroscopically hispid.

Young or small individuals of this species may be difficult to identify in the field, but larger individuals have a distinctive combination of growth form, color, and very hispid surface. All individuals of this form were correctly identified to species before verification with spicules and DNA sequencing, and no other samples were erroneously assigned to this species, so tentative field identifications are likely to be fairly reliable with experienced observers. Individuals from this species from Southern California were the focus of recent research on terpene biosynthesis in sponges (Wilson *et al*. 2023).

Order Clionaida

Family Clionaidae

***Cliona californiana* de Laubenfels, 1932**

Figure 1D.

**Synonymy**

*Cliona celata var. californiana* (de Laubenfels 1932)

*Pseudosuberites pseudos* (Dickinson 1945)

**Material examined.** IZC00048441, Inner Carmel Pinnacle (36.55852, −121.96820), 10–24 m, 9/22/21.

**Morphology.** Sampled in the alpha stage, this sponge was visible only as small golden papillae embedded within unoccupied barnacle tests. Breaking these barnacle tests revealed additional sponge tissue within, which was sampled for DNA extraction and spicule preparation.

**Skeleton.** Not investigated.

**Spicules.** Tylostyles: curved; styles slightly subterminal in nearly all cases. 175–257–305 x 5–9– 11 μm (n=34).

**Distribution and habitat.** Southern Alaska to Oaxaca, Mexico, intertidal to deep water (Austin 1985; Carballo *et al*. 2004; Lee *et al*. 2007).

**Remarks.** This sponge was sampled from near the type location for the species, the intertidal zone in Pacific Grove (Monterey Peninsula), and the genetic data from *cox1* and *28S* are the first available for the species. Closely related taxa in the Atlantic and South Pacific have been found to contain cryptic species (de Paula *et al*. 2012), so these new genetic data will be useful in future investigations of populations on the West Coast of North America.

Order Tetractinellida

Family Ancorinidae

***Stelletta clarella* de Laubenfels, 1930**

Figure 25

**Figure 25.**
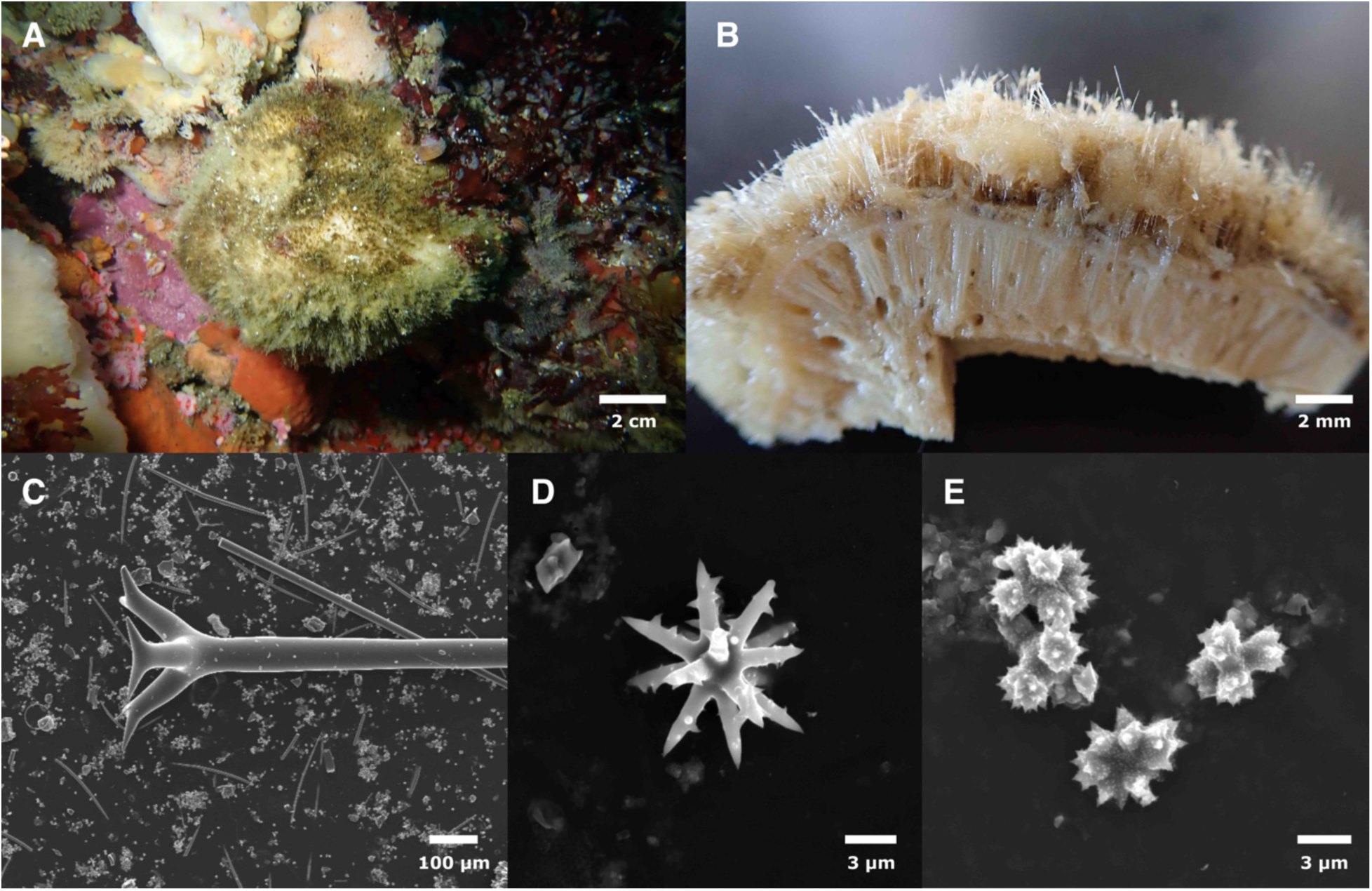
*Stelletta clarella*. A: Field photo. B: Cross-section showing spicule plush. C: Dichotriaene; small foreign styles and abundant asters visible in the background. D: Oxyaster. E: Spheroxyasters. All images from IZC00048471.

**Material examined.** IZC00048471, Inner Carmel Pinnacle (36.55852, −121.96820), 10–24 m, 9/22/21.

**Morphology.** Globular, 10 cm across, with 5 mm of spicule plush covering the outside and trapping copious debris. White alive and when preserved.

**Skeleton.** Not investigated.

**Spicules.** Oxeas, Dichotriaenes, Plagiotriaenes, oxyasters and spheroxyasters; abundant styles also present but presumed foreign.

Oxeas: 3881–4913–6118 x 15–35–63 μm (n=11).

Dichotriaenes: Shaft lengths 2444–4214–5549 x 54–71–91 μm (n=11); chord lengths 150–257– 335 μm (n=14).

Plagiotriaenes: Shaft lengths 2334–3139–3944 x 23–57–76 μm (n=2); chord lengths 106–194– 253 μm (n=6).

Oxyasters: 10–12–13 μm (n=9).

Spheroxyasters: Small and irregular, with minute spines. 4–5–7 μm (n=22).

Styles: Abundant but presumed to be foreign because they are not known from this species, and have not been seen in other samples examined by the authors. 162–187–216 x 4–5–6 μm (n=7).

**Distribution and habitat.** Common in Northern and Central California, from the intertidal to deep water. Occurs as far north as southern Alaska. Additional work is needed to determine if *S. clarella* is also found in Southern California (see Remarks). Not known from artificial substrates.

**Remarks.** *Stelletta clarella* was described from Pescadero Point, near the Carmel Pinnacles where this sample was collected, and is the only *Stelletta* known from Central California. A second species, *S. estrella* de Laubenfels 1930, was described from Southern California, but this species was proposed to be a junior synonym of *S. clarella* by Bakus and Green (1987). Here we follow Lee (2007) and propose that *S. estrella* may be a junior synonym, but more work is needed to be sure.

Class Calcarea

Order Leucosolenida

Family Amphoriscidae

*Leucilla nuttingi* (Urban, 1902)

Figures 1I & 24A

**Synonymy.**

*Rhabdodermella nuttingi* (de Laubenfels 1932; Urban 1902)

**Material examined.** IZC00048454 and IZC00048455, Inner Carmel Pinnacle, (36.55910, − 121.96630), 10–18 m, 8/10/21; IZC00048463, Inner Carmel Pinnacle (36.55852, −121.96820), 10–24 m, 9/22/21.

**Morphology.** Previously described as vase shaped or tubular, up to 5 cm long with a stalk up to 2 cm, widest in center and narrowing to apical oscule (Lee *et al*. 2007). Samples examined are consistent with this description. Large specimens (figure 1I) 3 cm in vase length with 8 mm stalk (likely truncated when collected), 1.2 cm at widest point, 2 mm at apical osculum. Small sponge (figure 24A) 3 mm in vase length, 1 mm at widest point, 0.5 diameter at apical osculum.

**Skeleton.** Not examined.

**Spicules.** Large oxeas, oxeas, triradiates and quadriradiates.

Large coronal oxeas: Only present in the small sample 1248–1775–1796 x 21–23–25 μm (n=4).

Oxeas: Tapering asymmetrically, sometimes with prominent serrations. Significantly longer in large samples. Large sample (IZC00048455): 161–196–279 x 1–2–3 μm (n=19), small sample (IZC00048456): 78–123–220 x 1–2–3 μm (n=19).

Triradiates and quadriradiates: In large sample, all triradiates were sagittal and large quadriradiates were common. In small samples, most triradiates were saggital but some were equilateral, quadriradiates less common. Significantly larger in large sample. Unpaired ray lengths, large sample (IZC00048455): 83–374–832 μm (n=114), small samples (IZC00048456): 77–225–639 μm (n=44).

**Distribution and habitat.** Known from the Aleutian Islands to Southern California, from the intertidal zone to 700 m (Austin 1985; Lee *et al*. 2007). Recent collections have found this species to be widespread and abundant in the rocky intertidal and subtidally at diving depths, on both natural and artificial reefs. Less commonly found on floating docks in marinas.

**Remarks.** We collected two clusters of large urns and one cluster of small urns from the Carmel Pinnacles, near the type location for *L. nuttingi*. We sequenced a 772 bp fragment of the *28S* locus from all three samples, and examined the spicules of one large and one small sample. The large and small samples had significant differences in their spicules, but were identical across 772 bp of the *28S* locus. Despite this variation in morphology, these samples all fit the descriptions of *L. nuttingi,* and no other species are known from the region that are similar (Lee *et al*. 2007). The Calcarea, however, have received scant attention in California since the work of Urban (1905), and additional work may reveal more species diversity.

## Conclusions

It is remarkable that, in only 90 minutes of search effort, we were able to collect 12 undescribed species at this one location. Half of these newly described species are thus far known only from locations in Carmel Bay, with four species known only from the Carmel Pinnacles SMR. The unique location and morphology of the Carmel Pinnacles likely combines a rare set of geological, biological, and oceanographic features to generate a unique habitat for sponges and other sessile filter-feeding invertebrates. Further work at this location –– especially at other depths –– would no doubt lead to additional discoveries.

These results illustrate how much remains to be learned about the sponges of California, and show that presence/absence surveys by divers are an effective way to bridge this gap. The apparently narrow distributions of some of the newly discovered species should also motivate future efforts to include under-studied taxa like sponges in conservation planning and monitoring.

## Acknowledgements

Logistical and collecting support was provided by Jean de Marignac (NOAA) and many members of the UCSB dive program, especially Christoph Pierre, Christian Orsini, and Clint Nelson. Christina Piotrowski graciously provided access to vouchers from the California Academy of Sciences. We would also like to thank the editor and two anonymous reviewers for their help in improving the paper. The scientific results and conclusions, as well as any views or opinions expressed herein, are those of the authors and do not necessarily reflect the views of NOAA or the Department of Commerce.

